# In Silico Investigations of Adaptive Therapy Using Two Cytotoxic or Two Cytostatic Drugs

**DOI:** 10.1101/2023.05.12.540626

**Authors:** Daniel K. Saha, Alexander R. A. Anderson, Luis Cisneros, Carlo C. Maley

## Abstract

While the dose modulation (DM) protocols (DM Cocktail Tandem, DM Ping-Pong Alternate Every Cycle, DM Ping-Pong on Progression) involves adjusting drug dosages when the tumor burden changes, the fixed-dose (FD) protocols involves administering a specific, constant dosage of the drug only when the tumor is growing (Dose-Skipping) or when the absolute tumor burden is above the baseline level until it reduces to a certain percentage of the baseline (Intermittent). Moreover, two different drugs can be administered simultaneously (cocktail), or the drugs can be switched such that only one drug is applied at a given time (ping-pong), either every cycle (ping-pong alternate every cycle) or when the tumor grows (ping-pong on progression). The dose modulation protocols work well when treated with two cytotoxic drugs, however, the ping-pong protocols (DM Ping-Pong Alternate Every Cycle, DM Ping-Pong on Progression, FD Ping-Pong Intermittent) work well when treated with two cytostatic drugs. In general, adaptive therapy, using either two cytotoxic or two cytostatic drugs works best under conditions of high competition, such as high fitness cost, high replacement rates, and high turnover, although treatment using two cytostatic drugs works best under low turnover in many cases. Adaptive therapy works best when drug dosages are changed as soon as a change in tumor burden is detected, and it is best to pause treatment sooner rather than later when the tumor is shrinking. Adaptive therapy works best when an intermediate level of drug dosage is used, and both treatment with too little or too much drug leads to poor survival outcome.

## Introduction

Previously, we have investigated multi-drug adaptive therapy protocols for treatment using two drugs ^1^. Because the drugs investigated worked by killing the cells, the anti-cancer drugs would be expected to have a cytotoxic mechanism of action, killing cells directly ^2^. Cytostatic drugs, on the other hand, work by inhibiting cellular division ^2^. An interesting question that arises is, would the survival outcome be any different for adaptive therapy using two cytostatic drugs? We have undertaken to answer this question by investigating seven different adaptive therapy protocols and standard treatment at maximum tolerated dose, for treatment using either two cytotoxic drugs, or two cytostatic drugs, under a wide variety of different scenarios of cell kinetics and treatment settings. Here we are asking how the results change with cytostatic drugs, and two additional protocols, also capping the dose to a specific percentage of the maximum tolerated dose.

Experimental studies on adaptive therapy have been carried out in mice models of cancer using a single drug ^3, 4^, both these studies demonstrating improved survival outcome with adaptive therapy over standard treatment. A clinical trial of advanced stage metastatic prostate cancer patients using intermittent therapy resulted in an increased time to progression compared to a contemporaneous cohort of patients ^5, 6^. Adaptive therapy leverages fitness cost of resistance, which is the fitness penalty incurred by resistant cells in the absence of the drug, in order to maintain long-term control over the tumor ^1, 4, 5, 7–18^

## Materials and Methods

We have extended the original model^1^ in the following ways: the definition of progression (see *Observation*), the implementation of cytostatic drugs (see *Cell Death* and *Cell Division*). The changes to the original model ^1^ are as follows:

In section 2.4.11 (Observation), the change that have been made to the definition of progression, the modified survival criterion being: At any point after therapy initiation, if the tumor burden equaled or exceeded 97% of the carrying capacity, or if the rolling average of the total number of resistant cells over a period of 500 time-steps equaled or exceeded 50% of the carrying capacity, then we consider progression has occurred.

In section 2.7.1 (Cell Death), for treatment with two cytotoxic drugs, nothing has changed. For treatment using two cytostatic drugs, the equation for probability of cell death is as follows: Probability of cell death for a particular cell type per hour=background death probability of that particular cell type per hour.

In section 2.7.2 (Cell Division), for treatment with two cytotoxic drugs, nothing has changed. For treatment with two cytostatic drugs, probability of cell division per hour=background division probability per hour-S1*[Drug1]\***Ψ**1-S2*[Drug2]\***Ψ**2, where S1 and S2 are the binary indicator variables for the cell’s sensitivity to drugs 1 and 2, respectively, such that a value of 1 indicates sensitivity, while a value of 0 indicates resistance to the particular drug. [Drug1] and [Drug 2] being the concentrations of those drugs (non-negative real values), and **Ψ**1 and **Ψ**2 being the drug potency of those drugs (non-negative real values), quantified as the probability of inhibition in cell division per unit drug concentration per hour.

In section 2.7.6 (Drug Protocols), we investigated two additional treatment protocols—FD Ping-Pong Dose-Skipping and FD Ping-Pong Intermittent—as well as changed the nomenclature as follows:

FD Cocktail Dose-Skipping: It is identical to FD Dose-Skipping/Drug Holiday, which administers both the drugs, Drug1 and 2, in a cocktail formulation.

FD Ping-Pong Dose-Skipping: It is similar to FD Cocktail Dose-Skipping, other than Drug 1 and Drug 2 are being switched every treatment cycle, with the response of a particular drug being evaluated based on the how the tumor responded to that same drug last time it was administered.

FD Cocktail Intermittent: It is identical to FD Intermittent, except that Drugs 1 and 2 are administered as a cocktail formulation at 100% of the MTD (previously I used 75% of the MTD).

FD Ping-Pong Intermittent: It is similar to FD Cocktail Intermittent, other than the drugs are switched every time the tumor climbs back to 100% or more of the baseline tumor burden at which treatment was initiated.

## Results

### Cytotoxic and Cytostatic Therapies

For treatment using 2 cytotoxic drugs, all the dose modulation protocols (DM Cocktail Tandem, DM Ping-Pong Alternate Every Cycle, DM Ping-Pong on Progression), as well as FD Ping-Pong Dose-Skipping with a relatively low effect size as measured by the hazard ratio work well (Fig. 1A, Table 1) works well. For treatment using 2 cytostatic drugs, all the ping-pong protocols work well (DM Ping-Pong Alternate Every Cycle, DM Ping-Pong on Progression, FD Ping-Pong Intermittent) except the dose-skipping (Fig. 1B, Table 1).

**Figure 1:**
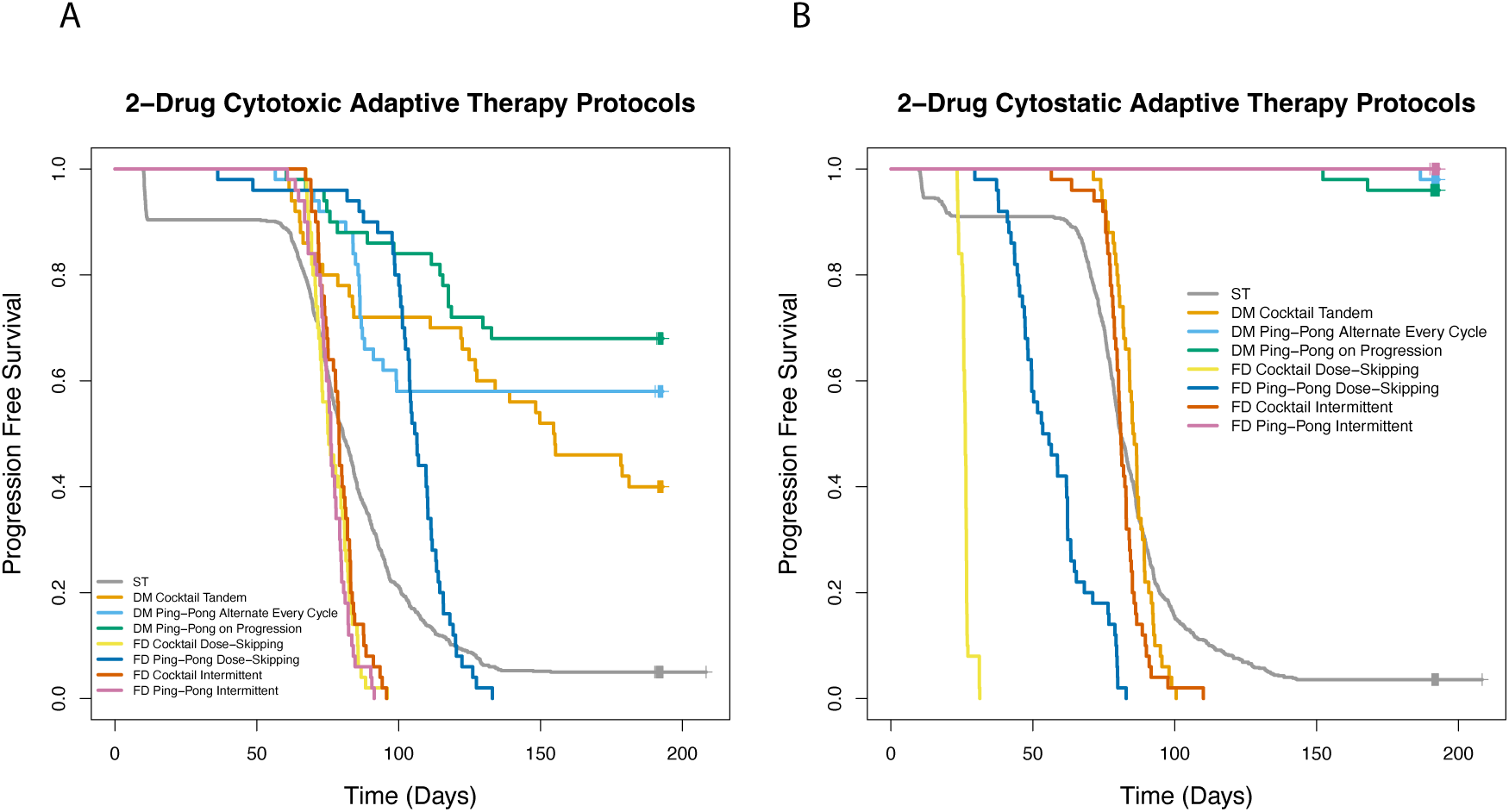
Adaptive therapy using two cytotoxic or two cytostatic drugs. Single-drug adaptive therapy protocols comparing standard treatment (ST) versus three different adaptive therapy protocols, dose modulation, dose-skipping, and intermittent using a single cytotoxic drug (Fig. 1A), or a single cytostatic drug (Fig. 1B).

**Table 1:**
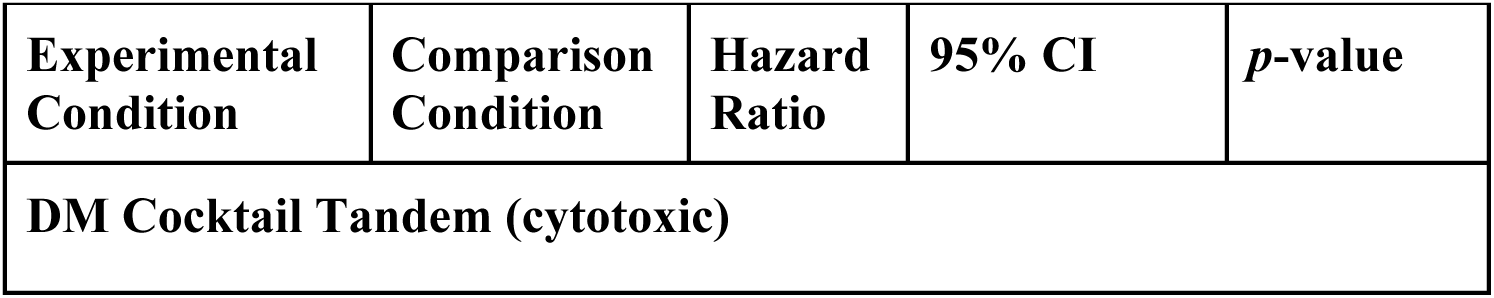
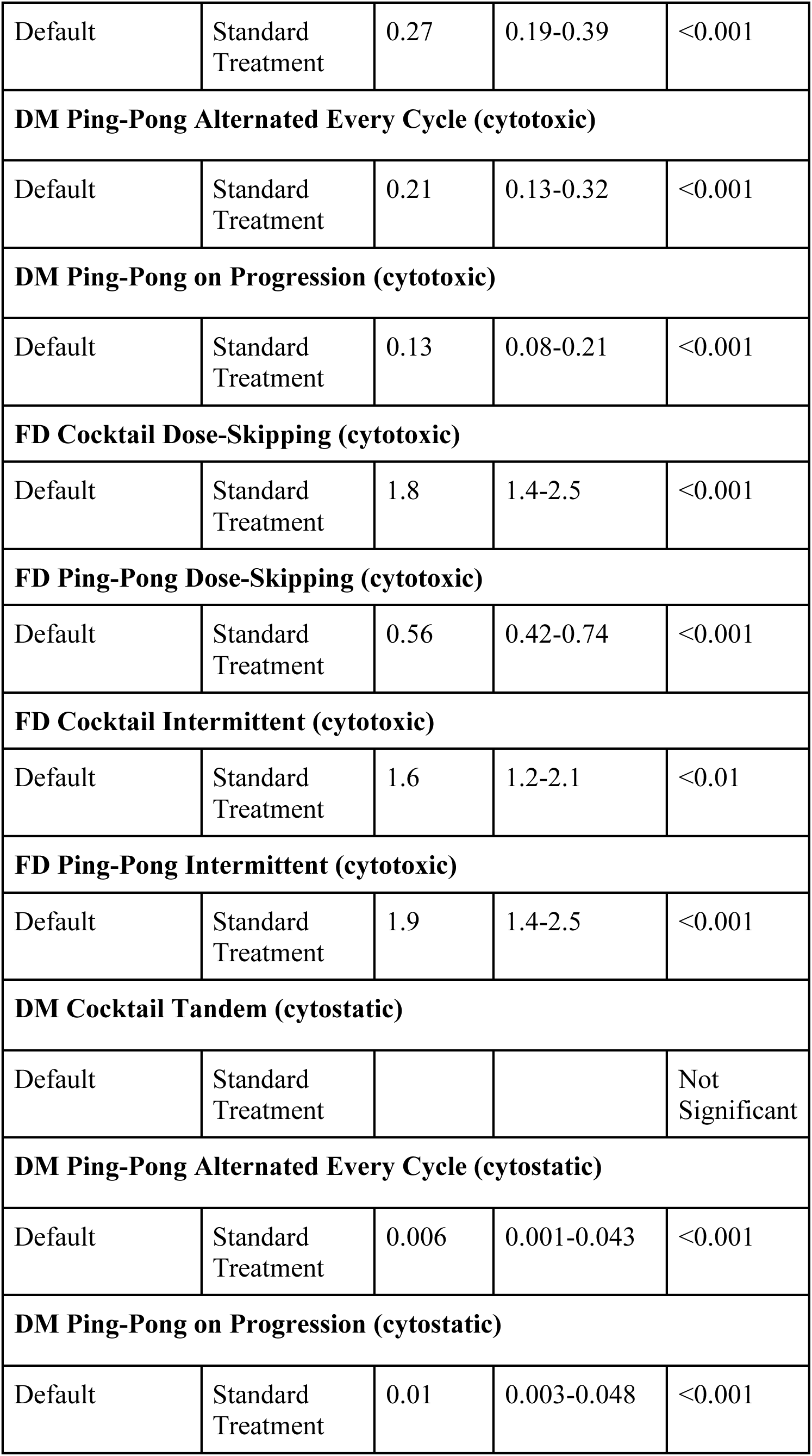
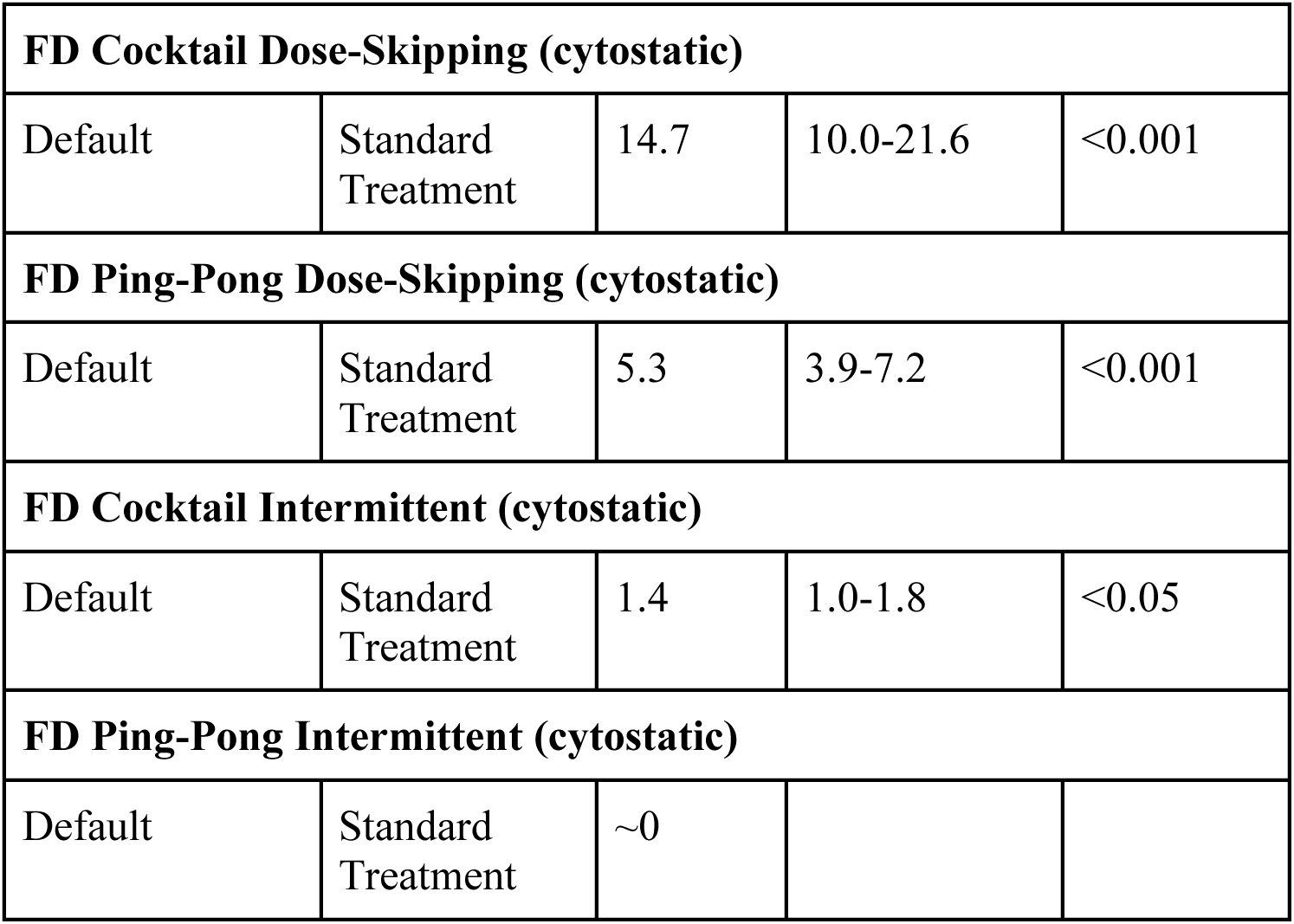
Adaptive therapy using two cytotoxic or two cytostatic drugs.

### The Effect of Fitness Costs of Resistance

For treatment using two cytotoxic drugs, all the dose-modulation protocols (DM Cocktail Tandem, DM Ping-Pong Alternate Every Cycle, DM Ping-Pong on Progression) work well under conditions of 5-fold or 7-fold fitness cost, and some work well even under conditions of 1.7-fold and 2.5-fold fitness cost, improving survival outcome relative to standard treatment (Fig. 2, Table 2); FD Ping-Pong works well under all fitness cost values (1.7-fold, 2.5-fold, 5-fold, 7-fold) tested (Table 2). For treatment using two cytostatic drugs, the ping-pong protocols (DM Ping-Pong Alternate Every Cycle, DM Ping-Pong on Progression, FD Ping-Pong Intermittent) work well under all values of fitness cost (Table 2); two exceptions can be noted to this general trend: FD Dose-Skipping under 5-fold or 7-fold fitness cost. In general, for treatment with either two cytotoxic or two cytostatic drugs, for the protocols that work, higher fitness cost values lead to improved survival outcome relative to low fitness cost as reflected in relatively low hazard ratios under conditions of higher fitness cost (Table 2).

**Figure 2.**
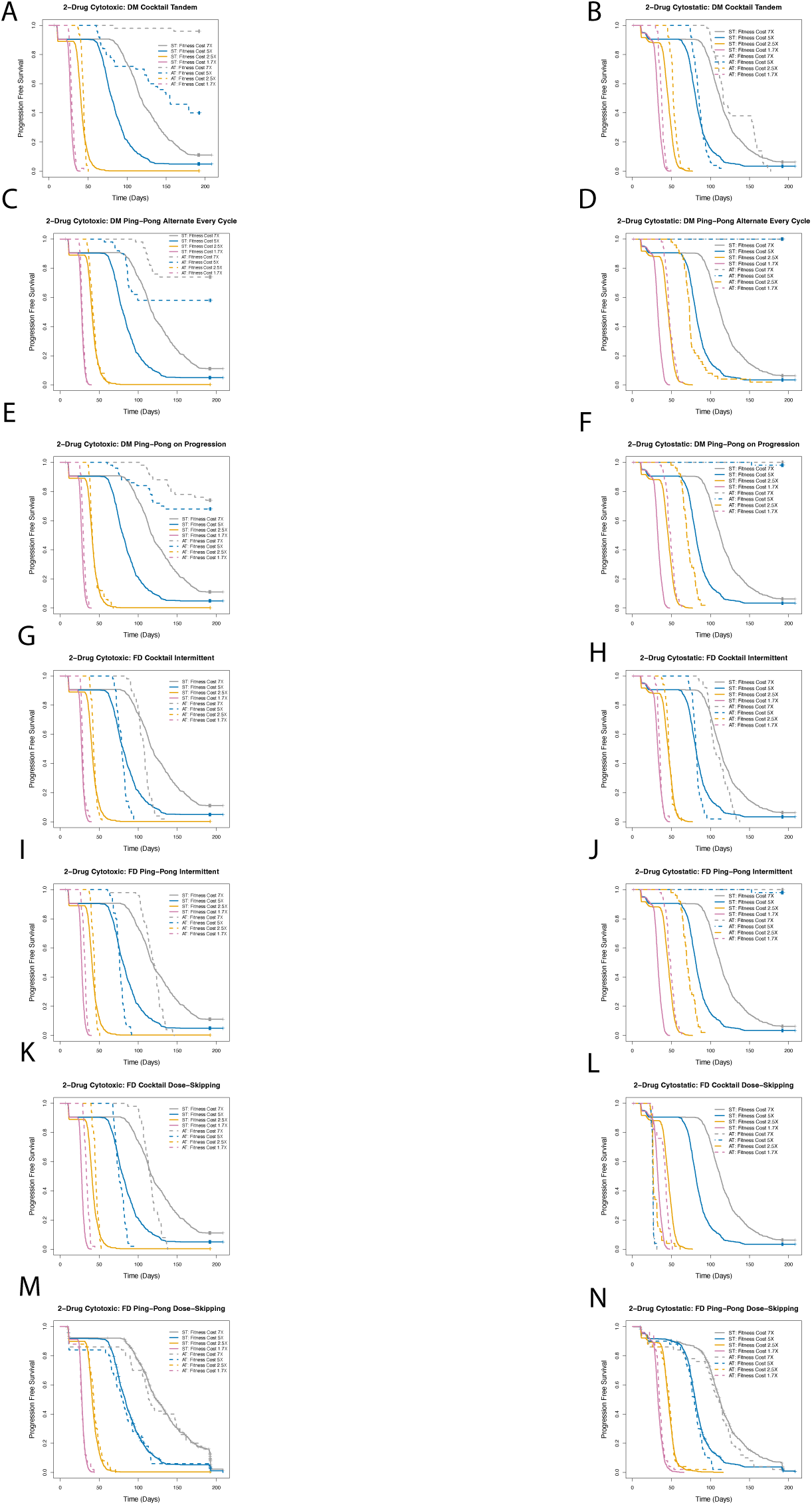
Effect of fitness cost parameter on the outcome of adaptive therapy using two cytotoxic or two cytostatic drugs. Survival outcome for treatment as per the dose modulation protocol (Fig. 2A, Fig. 2B), dose-skipping (Fig. 2C, Fig. 2D), or intermittent (Fig. 2E, Fig. 2F) under fitness cost of 1.7-fold, 2.5-fold, 5-fold, or 7-fold relative to standard treatment for treatment using either two cytotoxic (Fig. 2A, 2C, 2E), or two cytostatic drugs (Fig. 2B, 2D, 2F).

**Table 2:** Effect of fitness cost parameter on the outcome of adaptive therapy using two cytotoxic or two cytostatic drugs.

| Experimental Condition | Comparison Condition | Hazard Ratio | 95% CI | *p*-value |
| --- | --- | --- | --- | --- |
| **DM Cocktail Tandem (cytotoxic)** |  |  |  |  |
| 1.7-fold fitness cost | Standard Treatment | 0.62 | 0.46-0.82 | <0.01 |
| 2.5-fold fitness cost | Standard Treatment |  |  | Not Significant |
| 5-fold fitness cost | Standard Treatment | 0.27 | 0.19-0.39 | <0.001 |
| 7-fold fitness cost | Standard Treatment | 0.02 | 0.005-0.075 | <0.001 |
| **DM Ping-Pong Alternated Every Cycle (cytotoxic)** |  |  |  |  |
| 1.7-fold fitness cost | Standard Treatment |  |  | Not Significant |
| 2.5-fold fitness cost | Standard Treatment |  |  | Not Significant |
| 5-fold fitness cost | Standard Treatment | 0.21 | 0.13-0.32 | <0.001 |
| 7-fold fitness cost | Standard Treatment | 0.15 | 0.09-0.26 | <0.001 |
| **DM Ping-Pong on Progression (cytotoxic)** |  |  |  |  |
| 1.7-fold fitness cost | Standard Treatment | 0.57 | 0.43-0.76 | <0.001 |
| 2.5-fold fitness cost | Standard Treatment |  |  | Not Significant |
| 5-fold fitness cost | Standard Treatment | 0.13 | 0.08-0.21 | <0.001 |
| 7-fold fitness cost | Standard Treatment | 0.14 | 0.08-0.24 | <0.001 |
| **FD Cocktail Dose-Skipping (cytotoxic)** |  |  |  |  |
| 1.7-fold fitness cost | Standard Treatment | 0.23 | 0.17-0.32 | <0.001 |
| 2.5-fold fitness cost | Standard Treatment | 0.67 | 0.50-0.90 | <0.01 |
| 5-fold fitness cost | Standard Treatment | 1.8 | 1.4-2.5 | <0.001 |
| 7-fold fitness cost | Standard Treatment | 1.6 | 1.2-2.2 | <0.01 |
| **FD Ping-Pong Dose-Skipping (cytotoxic)** |  |  |  |  |
| 1.7-fold fitness cost | Standard Treatment | 0.18 | 0.13-0.25 | <0.001 |
| 2.5-fold fitness cost | Standard Treatment | 0.45 | 0.34-0.60 | <0.001 |
| 5-fold fitness cost | Standard Treatment | 0.56 | 0.42-0.74 | <0.001 |
| 7-fold fitness cost | Standard Treatment | 0.37 | 0.26-0.54 | <0.001 |
| **FD Cocktail Intermittent (cytotoxic)** |  |  |  |  |
| 1.7-fold fitness cost | Standard Treatment | 0.65 | 0.48-0.87 | <0.01 |
| 2.5-fold fitness cost | Standard Treatment |  |  | Not Significant |
| 5-fold fitness cost | Standard Treatment | 1.6 | 1.2-2.1 | <0.01 |
| 7-fold fitness cost | Standard Treatment | 2.3 | 1.7-3.1 | <0.001 |
| **FD Ping-Pong Intermittent (cytotoxic)** |  |  |  |  |
| 1.7-fold fitness cost | Standard Treatment | 0.43 | 0.32-0.57 | <0.001 |
| 2.5-fold fitness cost | Standard Treatment |  |  | Not Significant |
| 5-fold fitness cost | Standard Treatment | 1.9 | 1.4-2.5 | <0.001 |
| 7-fold fitness cost | Standard Treatment | 1.5 | 1.1-2.0 | <0.05 |
| **DM Cocktail Tandem (cytostatic)** |  |  |  |  |
| 1.7-fold fitness cost | Standard Treatment | 0.50 | 0.37-0.66 | <0.001 |
| 2.5-fold fitness cost | Standard Treatment | 0.47 | 0.35-0.63 | <0.001 |
| 5-fold fitness cost | Standard Treatment |  |  | Not Significant |
| 7-fold fitness cost | Standard Treatment |  |  | Not Significant |
| **DM Ping-Pong Alternated Every Cycle (cytostatic)** |  |  |  |  |
| 1.7-fold fitness cost | Standard Treatment | 0.07 | 0.05-0.11 | <0.001 |
| 2.5-fold fitness cost | Standard Treatment | 0.08 | 0.06-0.12 | <0.001 |
| 5-fold fitness cost | Standard Treatment | ~0 |  |  |
| 7-fold fitness cost | Standard Treatment | ~0 |  |  |
| **DM Ping-Pong on Progression (cytostatic)** |  |  |  |  |
| 1.7-fold fitness cost | Standard Treatment | 0.06 | 0.04-0.10 | <0.001 |
| 2.5-fold fitness cost | Standard Treatment | 0.10 | 0.07-0.14 | <0.001 |
| 5-fold fitness cost | Standard Treatment | 0.006 | 0.001-0.042 | <0.001 |
| 7-fold fitness cost | Standard Treatment | ~0 |  |  |
| **FD Cocktail Dose-Skipping (cytostatic)** |  |  |  |  |
| 1.7-fold fitness cost | Standard Treatment | 0.22 | 0.16-0.31 | <0.001 |
| 2.5-fold fitness cost | Standard Treatment | 4.0 | 2.9-5.4 | <0.001 |
| 5-fold fitness cost | Standard Treatment | 13.8 | 9.4-20.2 | <0.001 |
| 7-fold fitness cost | Standard Treatment | 13.8 | 9.4-20.2 | <0.001 |

| **FD Ping-Pong Dose-Skipping (cytostatic)** |  |  |  |  |
| --- | --- | --- | --- | --- |
| 1.7-fold fitness cost | Standard Treatment | 0.13 | 0.08-0.18 | <0.001 |
| 2.5-fold fitness cost | Standard Treatment | 0.35 | 0.25-0.48 | <0.001 |
| 5-fold fitness cost | Standard Treatment | 6.9 | 5.0-9.5 | <0.001 |
| 7-fold fitness cost | Standard Treatment | 13.6 | 9.3-19.8 | <0.001 |
| **FD Cocktail Intermittent (cytostatic)** |  |  |  |  |
| 1.7-fold fitness cost | Standard Treatment | 0.72 | 0.54-0.96 | <0.05 |
| 2.5-fold fitness cost | Standard Treatment |  |  | Not Significant |
| 5-fold fitness cost | Standard Treatment |  |  | Not Significant |
| 7-fold fitness cost | Standard Treatment | 1.6 | 1.2-2.1 | <0.01 |
| **FD Ping-Pong Intermittent (cytostatic)** |  |  |  |  |
| 1.7-fold fitness cost | Standard Treatment | 0.07 | 0.05-0.11 | <0.001 |
| 2.5-fold fitness cost | Standard Treatment | 0.10 | 0.07-0.15 | <0.001 |
| 5-fold fitness cost | Standard Treatment | 0.006 | 0.001-0.042 | <0.001 |
| 7-fold fitness cost | Standard Treatment | ~0 |  |  |

For treatment using two cytotoxic drugs, all the dose-modulation protocols (DM Cocktail Tandem, DM Ping-Pong Alternate Every Cycle, DM Ping-Pong on progression) as well as FD Ping-Pong Dose-Skipping work well under all conditions of replacement rate tested, that is, 0%, 50%, or 100% replacement. For treatment with two cytostatic drugs, the ping-pong protocols (DM Ping-Pong Alternate Every Cycle, DM Ping-Pong on Progression, FD Ping-Pong Intermittent) work well under all conditions of replacement tested here, with the exception of FD Ping-Pong Intermittent at 0% replacement where the effect was not significant. In general, when a protocol works, for either treatment using two cytotoxic or two cytostatic drugs, 100% replacement works best, followed by 50% replacement, and 0% replacement works worst, as reflected in the relatively low hazard ratios for higher replacement rates (Fig. 3, Table 3). Some protocols work so well (such as some of the ping-protocols using two cytostatic drugs) that the percent of replacement doesn’t matter as it was able to control the tumor under all conditions (Fig. 3D, 3F).

**Figure 3.**
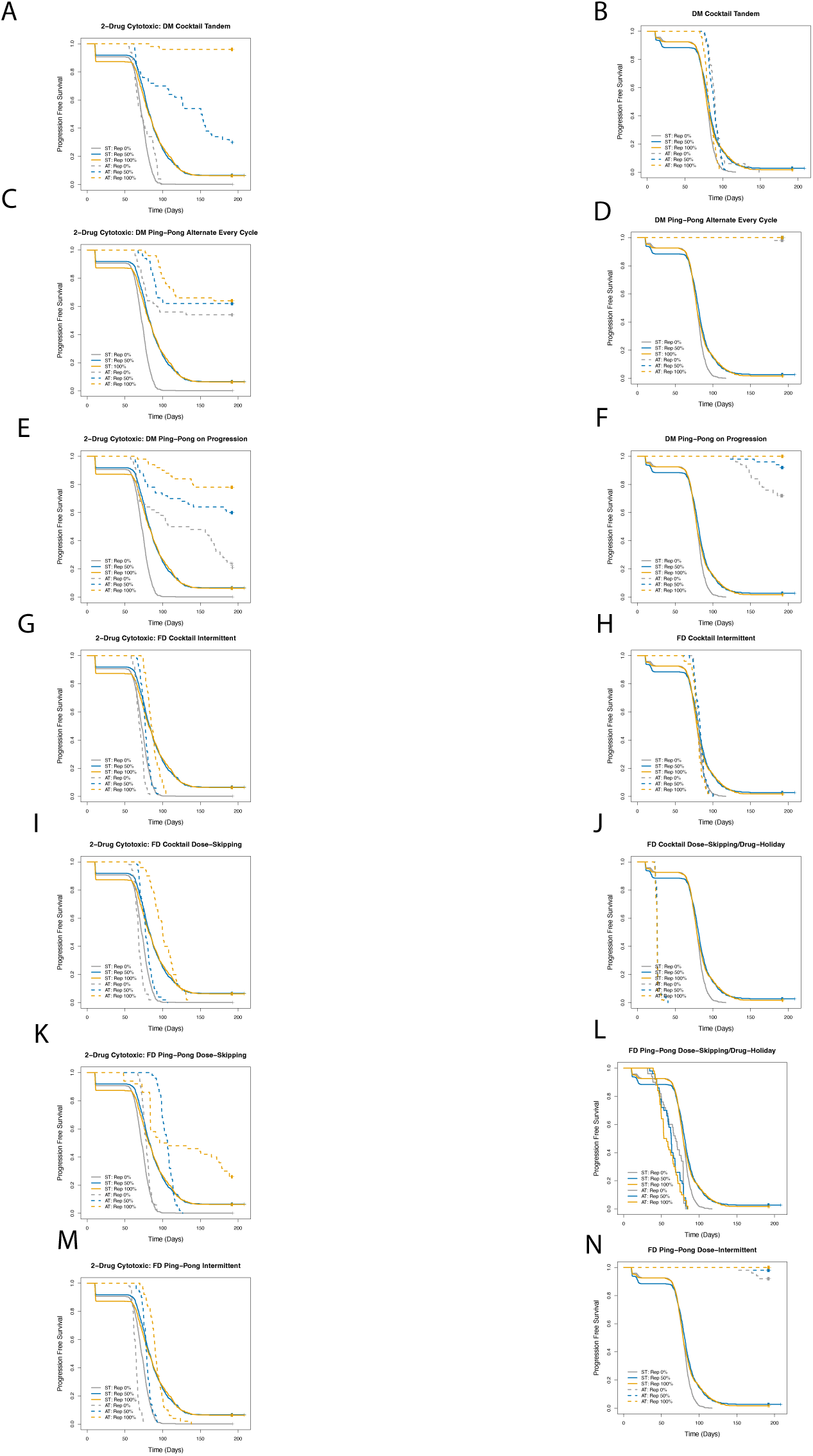
Effect of replacement parameter on outcome of adaptive therapy using two cytotoxic or two cytostatic drugs. Treatment as per the dose modulation protocol (Fig. 3A, 3B), dose-skipping protocol (Fig. 3C, Fig. 3D), or intermittent (Fig. 3E, Fig. 3F), relative to standard treatment under conditions of 0%, 50%, or 100% replacement using either a single cytotoxic (Fig. 3A,3C, 3E), or a single cytostatic drug (Fig. 3B, Fig. 3D, Fig. 3F).

**Table 3:** Effect of replacement parameter on outcome of adaptive therapy using two cytotoxic or two cytostatic drugs.

| Experimental Condition | Comparison Condition | Hazard Ratio | 95% CI | *p*-value |
| --- | --- | --- | --- | --- |
| **DM Cocktail Tandem (cytotoxic)** |  |  |  |  |
| 0% replacement | Standard Treatment | 0.64 | 0.48-0.86 | <0.01 |
| 50% replacement | Standard Treatment | 0.37 | 0.26-0.52 | <0.001 |
| 100% replacement | Standard Treatment | 0.01 | 0.004-0.060 | <0.001 |
| **DM Ping-Pong Alternated Every Cycle (cytotoxic)** |  |  |  |  |
| 0% replacement | Standard Treatment | 0.13 | 0.08-0.20 | <0.001 |
| 50% replacement | Standard Treatment | 0.19 | 0.12-0.30 | <0.001 |
| 100% replacement | Standard Treatment | 0.16 | 0.10-0.25 | <0.001 |
| **DM Ping-Pong on Progression (cytotoxic)** |  |  |  |  |
| 0% replacement | Standard Treatment | 0.16 | 0.11-0.24 | <0.001 |
| 50% replacement | Standard Treatment | 0.19 | 0.12-0.30 | <0.001 |
| 100% replacement | Standard Treatment | 0.09 | 0.05-0.16 | <0.001 |
| **FD Cocktail Dose-Skipping (cytotoxic)** |  |  |  |  |
| 0% replacement | Standard Treatment | 1.9 | 1.4-2.5 | <0.001 |
| 50% replacement | Standard Treatment | 1.8 | 1.3-2.4 | <0.001 |
| 100% replacement | Standard Treatment | 0.72 | 0.54-0.97 | <0.05 |
| **FD Ping-Pong Dose-Skipping (cytotoxic)** |  |  |  |  |
| 0% replacement | Standard Treatment | 0.70 | 0.52-0.93 | <0.05 |
| 50% replacement | Standard Treatment | 0.64 | 0.48-0.85 | <0.01 |
| 100% replacement | Standard Treatment | 0.44 | 0.31-0.61 | <0.001 |
| **FD Cocktail Intermittent (cytotoxic)** |  |  |  |  |
| 0% replacement | Standard Treatment | 1.7 | 1.2-2.2 | <0.001 |
| 50% replacement | Standard Treatment | 2.0 | 1.5-2.7 | <0.001 |
| 100% replacement | Standard Treatment |  |  | Not Significant |
| **FD Ping-Pong Intermittent (cytotoxic)** |  |  |  |  |
| 0% replacement | Standard Treatment | 3.3 | 2.4-4.4 | <0.001 |
| 50% replacement | Standard Treatment | 1.8 | 1.4-2.4 | <0.001 |
| 100% replacement | Standard Treatment |  |  | Not Significant |
| **DM Cocktail Tandem (cytostatic)** |  |  |  |  |
| 0% replacement | Standard Treatment | 0.41 | 0.30-0.55 | <0.001 |
| 50% replacement | Standard Treatment |  |  | Not Significant |
| 100% replacement | Standard Treatment |  |  | Not Significant |
| **DM Ping-Pong Alternated Every Cycle (cytostatic)** |  |  |  |  |
| 0% replacement | Standard Treatment | ~0 |  |  |
| 50% replacement | Standard Treatment | ~0 |  |  |
| 100% replacement | Standard Treatment | ~0 |  |  |
| **DM Ping-Pong on Progression (cytostatic)** |  |  |  |  |
| 0% replacement | Standard Treatment | ~0 |  |  |
| 50% replacement | Standard Treatment | 0.02 | 0.008-0.058 | <0.001 |
| 100% replacement | Standard Treatment | ~0 |  |  |
| **FD Cocktail Dose-Skipping (cytostatic)** |  |  |  |  |
| 0% replacement | Standard Treatment | 19.1 | 12.7-28.7 | <0.001 |
| 50% replacement | Standard Treatment | 10.5 | 7.3-15.1 | <0.001 |
| 100% replacement | Standard Treatment | 19.1 | 12.7-28.6 | <0.001 |
| **FD Ping-Pong Dose-Skipping (cytostatic)** |  |  |  |  |
| 0% replacement | Standard Treatment | 2.8 | 2.1-3.8 | <0.001 |
| 50% replacement | Standard Treatment | 3.9 | 2.9-5.3 | <0.001 |
| 100% replacement | Standard Treatment | 4.6 | 3.4-6.2 | <0.001 |
| **FD Cocktail Intermittent (cytostatic)** |  |  |  |  |
| 0% replacement | Standard Treatment |  |  | Not Significant |
| 50% replacement | Standard Treatment |  |  | Not Significant |
| 100% replacement | Standard Treatment |  |  | Not Significant |
| **FD Ping-Pong Intermittent (cytostatic)** |  |  |  |  |
| 0% replacement | Standard Treatment |  |  | Not Significant |
| 50% replacement | Standard Treatment | 0.01 | 0.001-0.039 | <0.001 |
| 100% replacement | Standard Treatment | ~0 |  |  |

For treatment using two cytotoxic drugs, the dose modulation protocols (DM Cocktail Tandem, DM Ping-Pong Alternate Every Cycle, DM Ping-Pong on Progression), as well as FD Ping-Pong Dose-Skipping works well under conditions of high turnover, some also working under low turnover conditions. In contrast, for treatment using two cytostatic drugs, the ping-pong protocols (DM Ping-Pong Alternate Every Cycle, DM Ping-Pong on Progression, FD Ping-Pong Intermittent) works better under low turnover conditions; however, as an exception, FD Ping-Pong Dose-Skipping works best under high turnover conditions (Fig. 4L, 4N, Table 4). In general, cytostatic drugs tend to work better when turnover is low, relative to when turnover is high, including the standard treatment at maximum tolerated dose, as indicated by the low hazard ratios under low turnover conditions (Table 4). For cytotoxic drugs, adaptive therapy works better in tumors with high turnover (Fig. 4).

**Figure 4.**
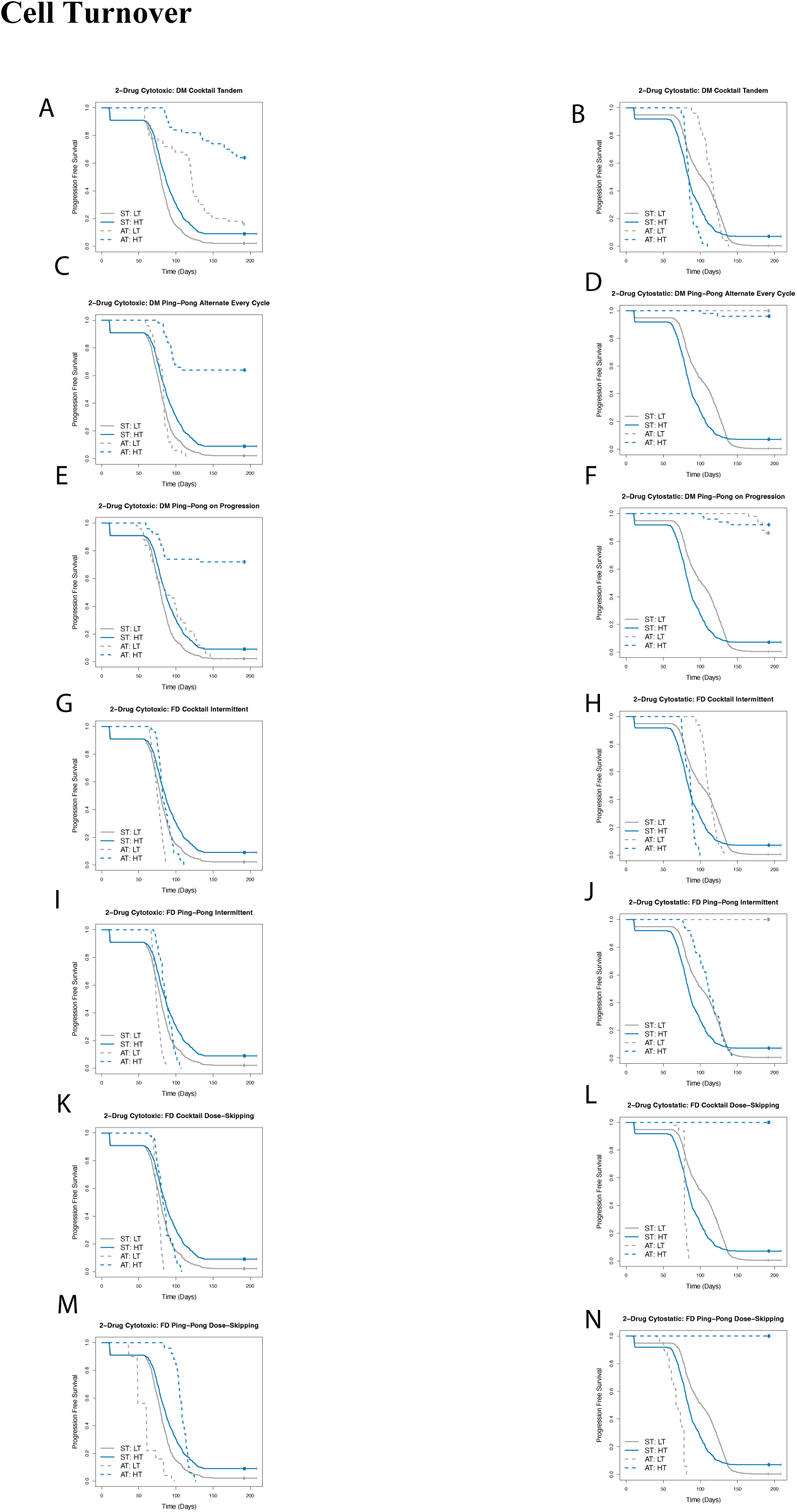
Effect of turnover on outcome of adaptive therapy using two cytotoxic or two cytostatic drugs. Survival outcome for treatment as per the dose modulation protocol (Fig. 4A, Fig. 4B), dose-skipping protocol (Fig. 4C, Fig. 4D), or intermittent (Fig. 4E, Fig. 4F) using a single cytotoxic (Fig. 4A, 4C, 4E) or a single cytostatic drug (Fig. 4B, 4D, 4F) under conditions of low turnover (LT) or high turnover (HT), relative to standard treatment under those conditions.

**Table 4:** Effect of turnover on outcome of adaptive therapy using two cytotoxic or two cytostatic drugs.

| Experimental Condition | Comparison Condition | Hazard Ratio | 95% CI | p-value |
| --- | --- | --- | --- | --- |
| **DM Cocktail Tandem (cytotoxic)** |  |  |  |  |
| Low Turnover | Standard Treatment | 0.34 | 0.25-0.47 | <0.001 |
| High Turnover | Standard Treatment | 0.17 | 0.10-0.27 | <0.001 |

| **DM Ping-Pong Alternated Every Cycle (cytotoxic)** |  |  |  |  |
| --- | --- | --- | --- | --- |
| Low Turnover | Standard Treatment |  |  | Not Significant |
| High Turnover | Standard Treatment | 0.20 | 0.12-0.31 | <0.001 |
| **DM Ping-Pong on Progression (cytotoxic)** |  |  |  |  |
| Low Turnover | Standard Treatment | 0.67 | 0.50-0.89 | <0.01 |
| High Turnover | Standard Treatment | 0.15 | 0.09-0.26 | <0.001 |
| **FD Cocktail Dose-Skipping (cytotoxic)** |  |  |  |  |
| Low Turnover | Standard Treatment | 1.9 | 1.4-2.6 | <0.001 |
| High Turnover | Standard Treatment | 1.5 | 1.1-2.0 | <0.01 |
| **FD Ping-Pong Dose-Skipping (cytotoxic)** |  |  |  |  |
| Low Turnover | Standard Treatment | 3.4 | 2.6-4.6 | <0.001 |
| High Turnover | Standard Treatment | 0.68 | 0.51-0.90 | <0.01 |
| **FD Cocktail Intermittent (cytotoxic)** |  |  |  |  |
| Low Turnover | Standard Treatment | 1.8 | 1.4-2.5 | <0.001 |
| High Turnover | Standard Treatment | 1.5 | 1.1-2.0 | <0.01 |
| **FD Ping-Pong Intermittent (cytotoxic)** |  |  |  |  |
| Low Turnover | Standard Treatment | 2.0 | 1.5-2.7 | <0.001 |
| High Turnover | Standard Treatment | 1.3 | 1.0-1.8 | <0.05 |

| **DM Cocktail Tandem (cytostatic)** |  |  |  |  |
| --- | --- | --- | --- | --- |
| Low Turnover | Standard Treatment |  |  | Not Significant |
| High Turnover | Standard Treatment |  |  | Not Significant |
| **DM Ping-Pong Alternated Every Cycle (cytostatic)** |  |  |  |  |
| Low Turnover | Standard Treatment | ~0 |  |  |
| High Turnover | Standard Treatment | 0.02 | 0.004-0.062 | <0.001 |
| **DM Ping-Pong on Progression (cytostatic)** |  |  |  |  |
| Low Turnover | Standard Treatment | 0.02 | 0.01-0.05 | <0.001 |
| High Turnover | Standard Treatment | 0.03 | 0.01-0.08 | <0.001 |
| **FD Cocktail Dose-Skipping (cytostatic)** |  |  |  |  |
| Low Turnover | Standard Treatment | 4.3 | 3.1-5.9 | <0.001 |
| High Turnover | Standard Treatment | ~0 |  |  |
| **FD Ping-Pong Dose-Skipping (cytostatic)** |  |  |  |  |
| Low Turnover | Standard Treatment | 8.1 | 5.9-11.2 | <0.001 |
| High Turnover | Standard Treatment | ~0 |  |  |
| **FD Cocktail Intermittent (cytostatic)** |  |  |  |  |
| Low Turnover | Standard Treatment |  |  | Not Significant |
| High Turnover | Standard Treatment |  |  | Not Significant |

| FD Ping-Pong Intermittent (cytostatic) |  |  |  |  |
| --- | --- | --- | --- | --- |
| Low Turnover | Standard Treatment | ~0 |  |  |
| High Turnover | Standard Treatment | 0.57 | 0.43-0.76 | <0.001 |

### When to Adjust the Dose of the Drug

For treatment using two cytotoxic drugs, all the dose-modulation protocols (DM Cocktail Tandem, DM Ping-Pong Alternate Every Cycle, DM Ping-Pong on Progression) works well when Delta Tumor=5%, 10%, or 20% (Fig. 5, Table 5). For treatment using two cytostatic drugs, the ping-pong protocols (DM Ping-Pong Alternate Every Cycle, DM Ping-Pong on Progression) work well under Delta Tumor=5%, 10%, 20%, or 40% (Fig. 5, Table 5). In general, for both cytotoxic and cytostatic drugs changing the dose when the smallest change in tumor burden is detected works best (e.g., Delta Dose=5%) (Fig. 5, Table 5). In fact, with a delta dose of 5%, for the first time we observe that DM Cocktail Tandem with cytostatic drugs works well (Fig. 5B, Table 5).

**Figure 5.**
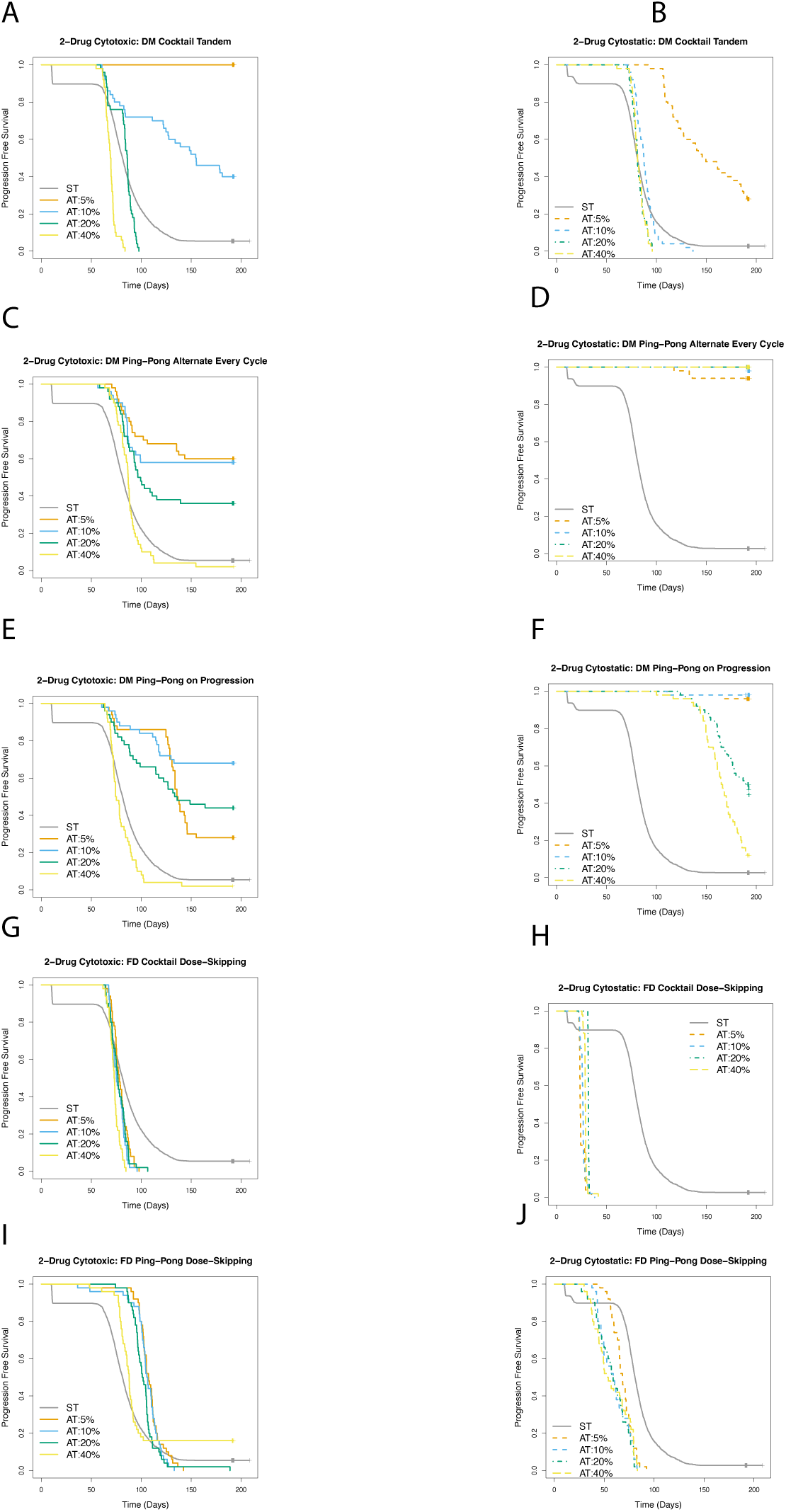
Effect of the delta tumor parameter on determining the outcome of adaptive therapy using two cytotoxic or two cytostatic drugs. Survival outcome comparing dose modulation treatment protocol with Delta Tumor=5%, 10%, 20%, or 40% using a single cytotoxic (Fig. 5A), or a single cytostatic drug (Fig. 5B) relative to standard treatment. Survival outcome comparing dose-skipping treatment protocol with Delta Tumor=5%, 10%, 20%, or 40% using a single cytotoxic (Fig. 5C), or a single cytostatic drug (Fig. 5D).

**Table 5:** Effect of the delta tumor parameter on determining the outcome of adaptive therapy using two cytotoxic or two cytostatic drugs.

| Experimental Condition | Comparison Condition | Hazard Ratio | 95% CI | *p*-value |
| --- | --- | --- | --- | --- |
| **DM Cocktail Tandem (cytotoxic)** |  |  |  |  |
| Delta Tumor=5% | Standard Treatment | ~0 |  |  |
| Delta Tumor=10% | Standard Treatment | 0.28 | 0.20-0.40 | <0.001 |
| Delta<br>Tumor=20% | Standard<br>Treatment |  |  | Not<br>Significant |
| Delta<br>Tumor=40% | Standard<br>Treatment | 3.4 | 2.6-4.6 | <0.001 |
| **DM Ping-Pong Alternated Every Cycle (cytotoxic)** |  |  |  |  |
| Delta<br>Tumor=5% | Standard<br>Treatment | 0.18 | 0.11-0.28 | <0.001 |
| Delta<br>Tumor=10% | Standard<br>Treatment | 0.21 | 0.13-0.32 | <0.001 |
| Delta<br>Tumor=20% | Standard<br>Treatment | 0.37 | 0.26-0.53 | <0.001 |
| Delta<br>Tumor=40% | Standard<br>Treatment |  |  | Not<br>Significant |
| **DM Ping-Pong on Progression (cytotoxic)** |  |  |  |  |
| Delta<br>Tumor=5% | Standard<br>Treatment | 0.29 | 0.21-0.41 | <0.001 |
| Delta<br>Tumor=10% | Standard<br>Treatment | 0.13 | 0.08-0.22 | <0.001 |
| Delta<br>Tumor=20% | Standard<br>Treatment | 0.27 | 0.19-0.40 | <0.001 |
| Delta<br>Tumor=40% | Standard<br>Treatment | 1.4 | 1.0-1.8 | <0.05 |
| **FD Cocktail Dose-Skipping (cytotoxic)** |  |  |  |  |
| Delta<br>Tumor=5% | Standard<br>Treatment | 1.6 | 1.2-2.1 | <0.01 |
| Delta<br>Tumor=10% | Standard<br>Treatment | 1.9 | 1.4-2.5 | <0.001 |
| Delta<br>Tumor=20% | Standard<br>Treatment | 1.7 | 1.3-2.3 | <0.001 |
| Delta<br>Tumor=40% | Standard<br>Treatment | 2.5 | 1.9-3.3 | <0.001 |

| **FD Ping-Pong Dose-Skipping (cytotoxic)** |  |  |  |  |
| --- | --- | --- | --- | --- |
| Delta Tumor=5% | Standard Treatment | 0.55 | 0.42-0.73 | <0.001 |
| Delta Tumor=10% | Standard Treatment | 0.58 | 0.44-0.77 | <0.001 |
| Delta Tumor=20% | Standard Treatment | 0.63 | 0.48-0.84 | <0.01 |
| Delta Tumor=40% | Standard Treatment | 0.70 | 0.52-0.95 | <0.05 |
| **DM Cocktail Tandem (cytostatic)** |  |  |  |  |
| Delta Tumor=5% | Standard Treatment | 0.22 | 0.16-0.31 | <0.001 |
| Delta Tumor=10% | Standard Treatment |  |  | Not Significant |
| Delta Tumor=20% | Standard Treatment |  |  | Not Significant |
| Delta Tumor=40% | Standard Treatment |  |  | Not Significant |
| **DM Ping-Pong Alternated Every Cycle (cytostatic)** |  |  |  |  |
| Delta Tumor=5% | Standard Treatment | 0.02 | 0.01-0.05 | <0.001 |
| Delta Tumor=10% | Standard Treatment | 0.01 | 0.001-0.039 | <0.001 |
| Delta Tumor=20% | Standard Treatment | ~0 |  |  |
| Delta Tumor=40% | Standard Treatment | ~0 |  |  |
| **DM Ping-Pong on Progression (cytostatic)** |  |  |  |  |
| Delta Tumor=5% | Standard Treatment | 0.01 | 0.003-0.045 | <0.001 |
| Delta<br>Tumor=10% | Standard<br>Treatment | 0.01 | 0.001-0.040 | <0.001 |
| Delta<br>Tumor=20% | Standard<br>Treatment | 0.14 | 0.09-0.20 | <0.001 |
| Delta<br>Tumor=40% | Standard<br>Treatment | 0.24 | 0.17-0.32 | <0.001 |
| **FD Cocktail Dose-Skipping (cytostatic)** |  |  |  |  |
| Delta<br>Tumor=5% | Standard<br>Treatment | 10.4 | 7.6-14.2 | <0.001 |
| Delta<br>Tumor=10% | Standard<br>Treatment | 10.4 | 7.6-14.2 | <0.001 |
| Delta<br>Tumor=20% | Standard<br>Treatment | 10.4 | 7.6-14.2 | <0.001 |
| Delta<br>Tumor=40% | Standard<br>Treatment | 10.4 | 7.6-14.2 | <0.001 |
| **FD Ping-Pong Dose-Skipping (cytostatic)** |  |  |  |  |
| Delta<br>Tumor=5% | Standard<br>Treatment | 3.3 | 2.5-4.4 | <0.001 |
| Delta<br>Tumor=10% | Standard<br>Treatment | 4.2 | 3.2-5.6 | <0.001 |
| Delta<br>Tumor=20% | Standard<br>Treatment | 4.4 | 3.3-5.9 | <0.001 |
| Delta<br>Tumor=40% | Standard<br>Treatment | 3.7 | 2.8-5.0 | <0.001 |

### How much to change the dose for the dose modulation protocols

For treatment using two cytotoxic drugs, all the dose modulation protocols (DM Cocktail Tandem, DM Ping-Pong Alternate Every Cycle, and DM Ping-Pong on Progression) work well when Delta Dose=25%, 50%, or 75%; with the exception of DM Cocktail Tandem with Delta Dose=25%. For treatment using two cytostatic drugs, the ping-pong protocols (DM Ping-Pong Alternate Every Cycle, DM Ping-Pong on Progression) works well when delta dose=25%, 50%, or 75%. In general, the larger delta doses work best, for both cytotoxic and cytostatic drugs (Fig. 6), with the exception of DM Ping-Pong on Progression with cytotoxic drugs (Fig. 6E, Table 6).

**Figure 6.**
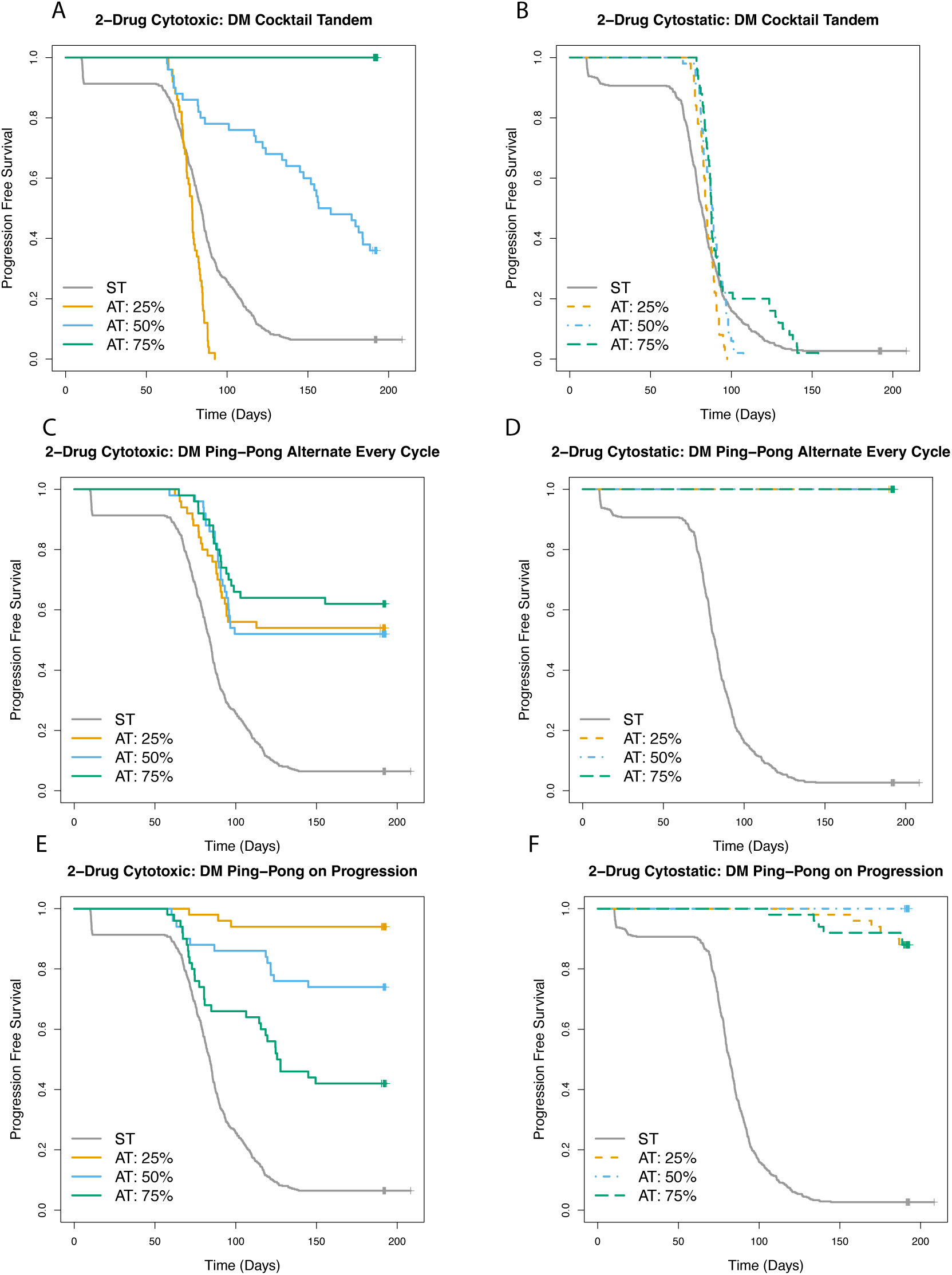
Effect of the delta dose parameter on determining the outcome of dose modulation adaptive therapy using two cytotoxic or two cytostatic drugs. Survival outcome for treatment as per the dose modulation protocol with Delta Dose=25%, 50%, or 75% relative to standard treatment using a single cytotoxic drug (Fig. 6A), or a single cytostatic drug (Fig. 6B).

**Table 6:** Effect of the delta dose parameter on determining the outcome of dose modulation adaptive therapy using two cytotoxic or two cytostatic drugs.

| Experimental Condition | Comparison Condition | Hazard Ratio | 95% CI | p-value |
| --- | --- | --- | --- | --- |
| **DM Cocktail Tandem (cytotoxic)** |  |  |  |  |
| Delta Dose=25% | Standard Treatment | 2.0 | 1.5-2.7 | <0.001 |
| Delta Dose=50% | Standard Treatment | 0.29 | 0.20-0.41 | <0.001 |
| Delta Dose=75% | Standard Treatment | ~0 |  |  |
| **DM Ping-Pong Alternated Every Cycle (cytotoxic)** |  |  |  |  |
| Delta Dose=25% | Standard Treatment | 0.26 | 0.17-0.40 | <0.001 |
| Delta<br>Dose=50% | Standard<br>Treatment | 0.26 | 0.17-0.40 | <0.001 |
| Delta<br>Dose=75% | Standard<br>Treatment | 0.19 | 0.12-0.30 | <0.001 |
| **DM Ping-Pong on Progression (cytotoxic)** |  |  |  |  |
| Delta<br>Dose=25% | Standard<br>Treatment | 0.02 | 0.01-0.07 | <0.001 |
| Delta<br>Dose=50% | Standard<br>Treatment | 0.11 | 0.06-0.19 | <0.001 |
| Delta<br>Dose=75% | Standard<br>Treatment | 0.32 | 0.22-0.47 | <0.001 |
| **DM Cocktail Tandem (cytostatic)** |  |  |  |  |
| Delta<br>Dose=25% | Standard<br>Treatment |  |  | Not<br>Significant |
| Delta<br>Dose=50% | Standard<br>Treatment |  |  | Not<br>Significant |
| Delta<br>Dose=75% | Standard<br>Treatment | 0.69 | 0.51-0.92 | <0.05 |
| **DM Ping-Pong Alternated Every Cycle (cytostatic)** |  |  |  |  |
| Delta<br>Dose=25% | Standard<br>Treatment | ~0 |  |  |
| Delta<br>Dose=50% | Standard<br>Treatment | ~0 |  |  |
| Delta<br>Dose=75% | Standard<br>Treatment | ~0 |  |  |
| **DM Ping-Pong on Progression (cytostatic)** |  |  |  |  |
| Delta<br>Dose=25% | Standard<br>Treatment | 0.03 | 0.01-0.07 | <0.001 |
| Delta<br>Dose=50% | Standard<br>Treatment | ~0 |  |  |
| Delta<br>Dose=75% | Standard<br>Treatment | 0.03 | 0.01-0.07 | <0.001 |

### Effects of pausing treatment when tumor burden falls below a certain level

For treatment using two cytotoxic or two cytostatic drugs, the dose-modulation protocols (DM Cocktail Tandem, DM Ping-Pong Alternate Every Cycle, DM Ping-Pong on Progression) work best when treatment is paused sooner than later when the tumor is responding, that is, triggering a treatment vacation when the tumor shrinks by 20%, for example, works better than waiting for the tumor to shrink by 50%, or 90% (Fig. 7, Table 7). The only exception is DM Ping-Pong on Progression (Fig. 7E), where pausing treatment at 50% is working better than 20%. A similar effect is observed for treatment using two cytotoxic (Fig. 7G, Fig 7I, Table 7) or two cytostatic drugs (Fig. 7H, 7J, Table 7) as per the intermittent protocol, where using a lower threshold for pausing treatment works better, however, the effect size is not strong for treatment using two cytotoxic drugs. In general, stopping treatment sooner than later, when the tumor is responding works well, for both treatment using two cytotoxic or two cytostatic drugs.

**Figure 7.**
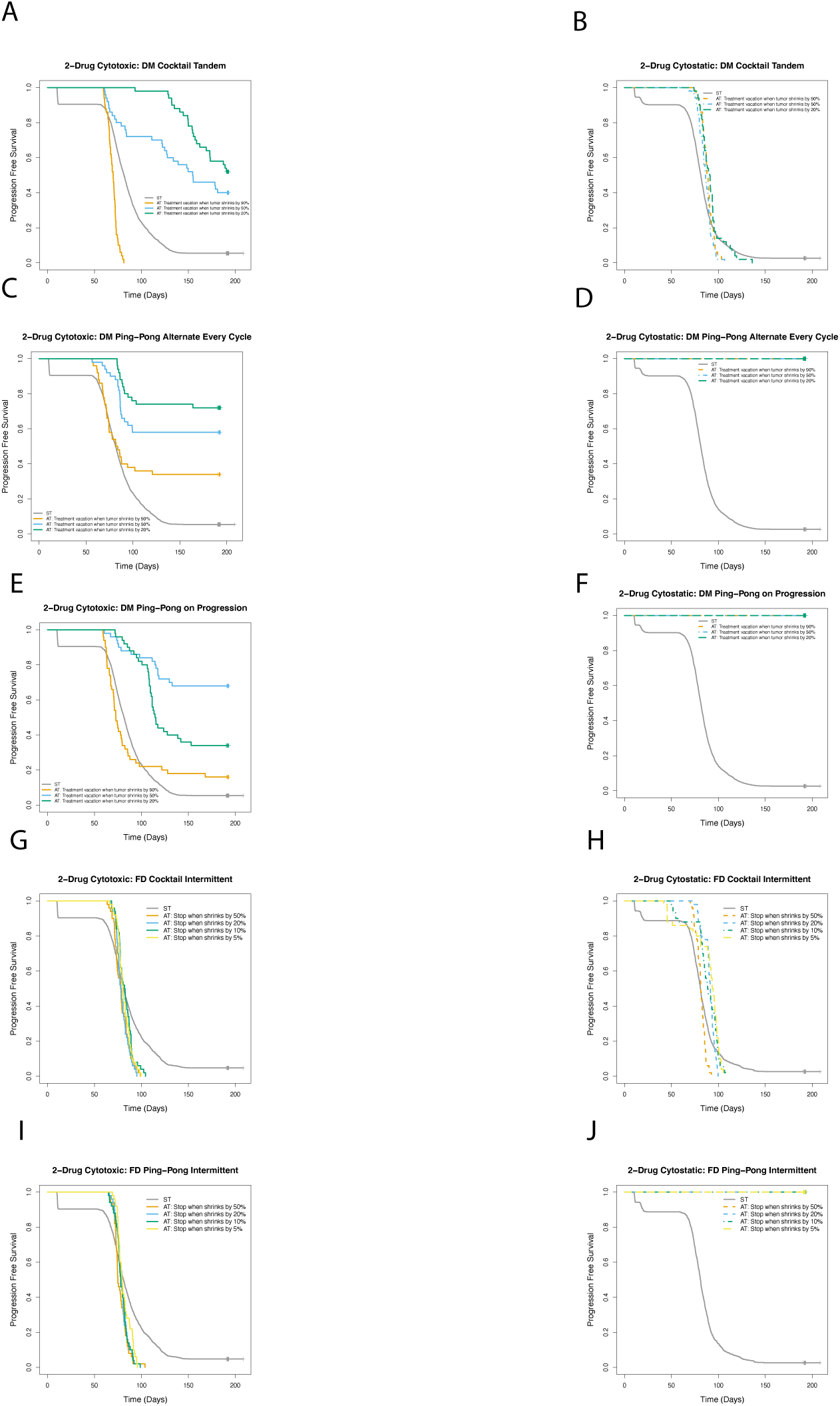
Effect of stopping treatment when tumor burden falls below a certain level for treatment using two cytotoxic or two cytostatic drugs. For treatment as per the intermittent protocol using either a single cytotoxic (Fig. 7A), or a single cytostatic drug (Fig. 7B), the threshold for stopping treatment was varied as the tumor shrinks by 5%, 10%, 20%, or 50% of the pre-treatment baseline. Survival outcome for treatment using a single cytotoxic drug (Fig. 7C), or a single cytostatic drug (Fig. 7D) as the trigger for treatment vacation is when the tumor shrinks by 20%, 50%, or 90%.

**Table 7:**
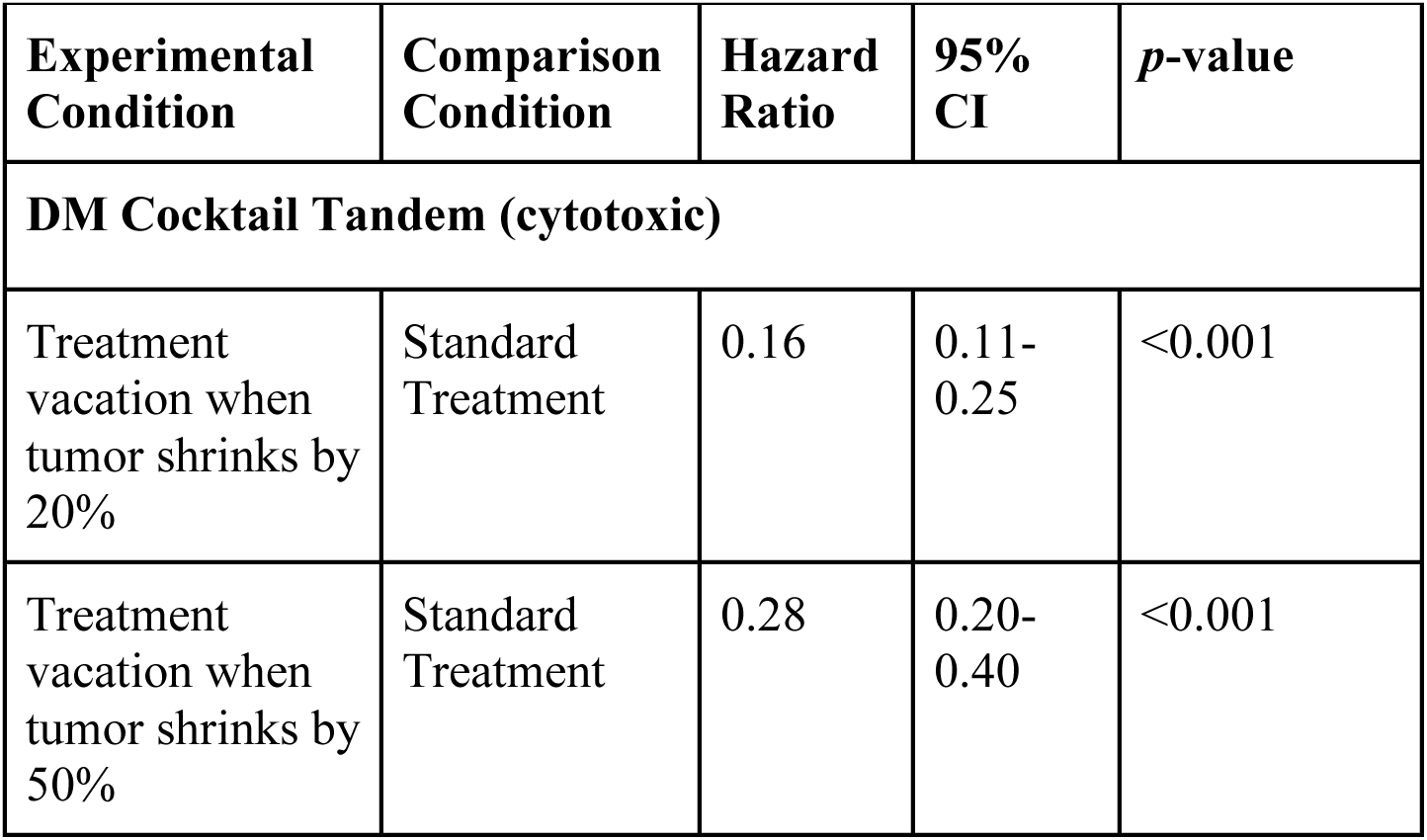
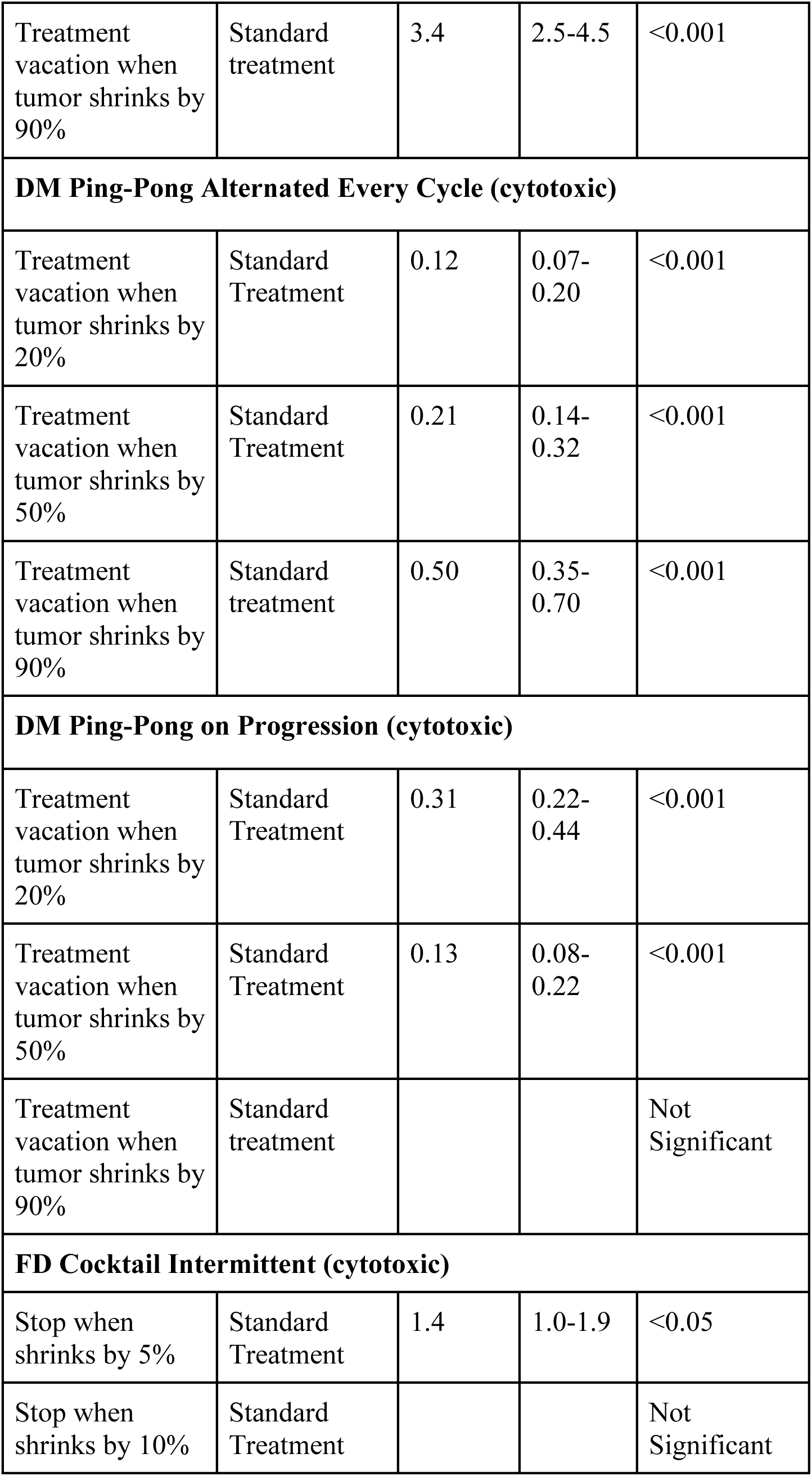
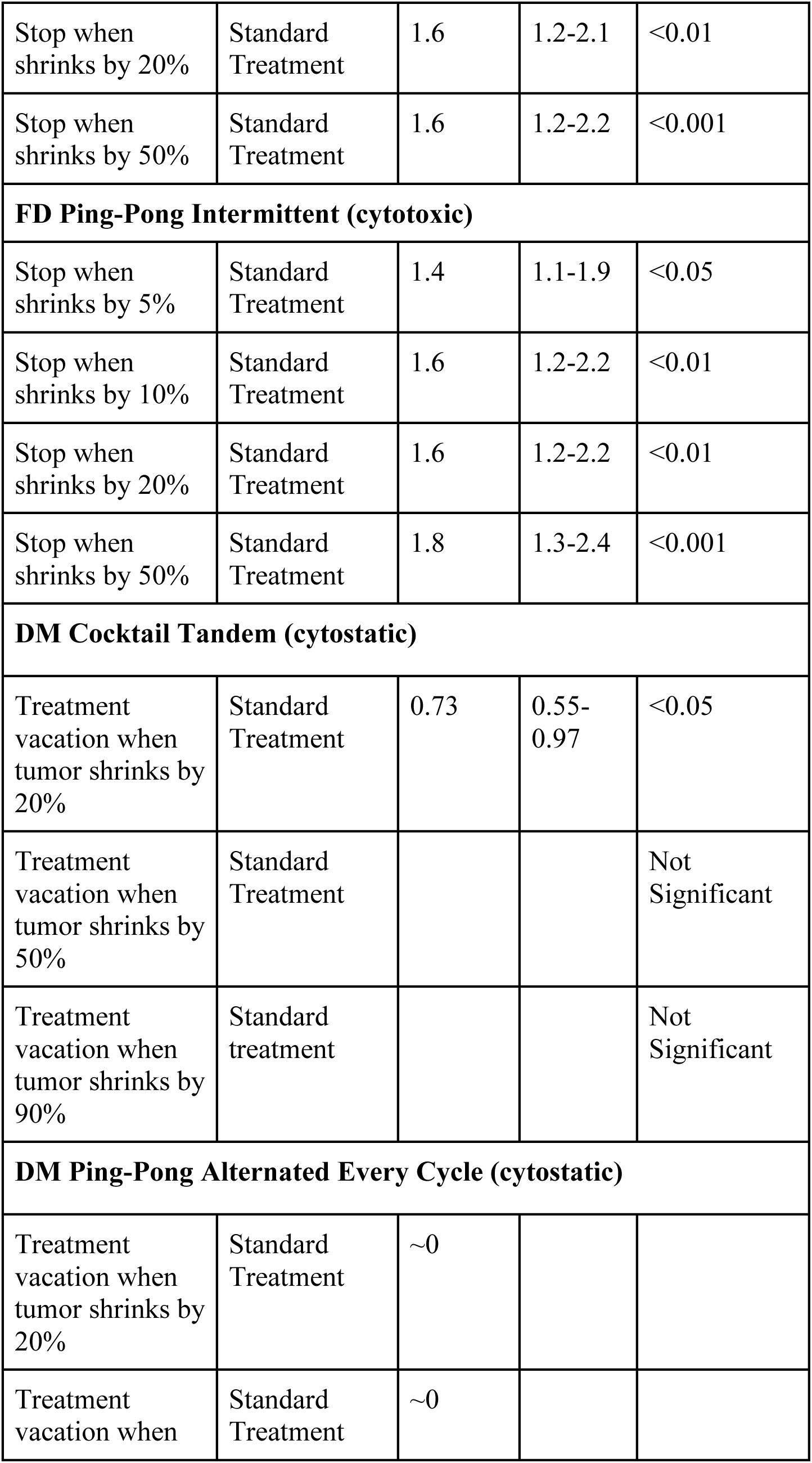
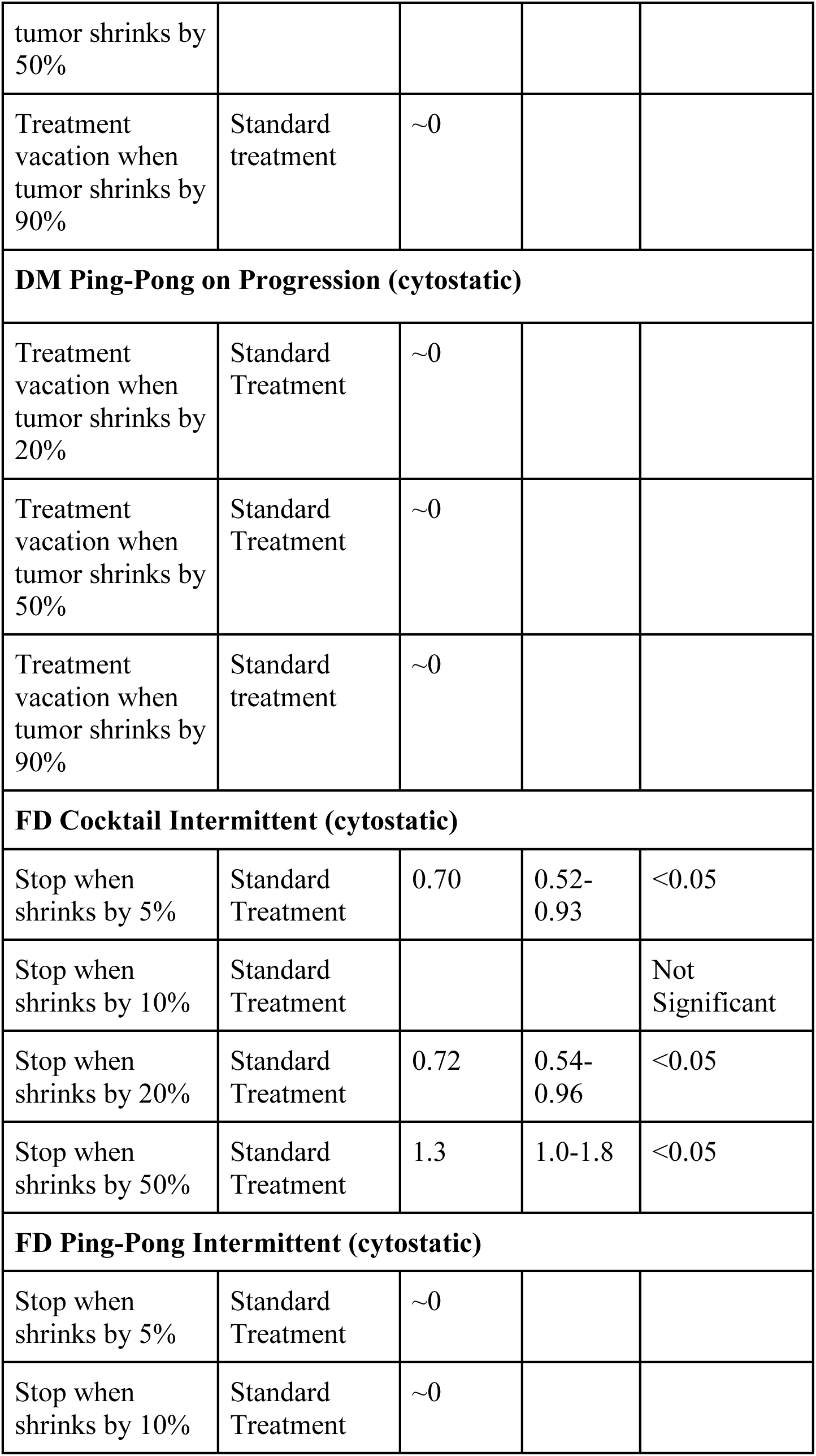
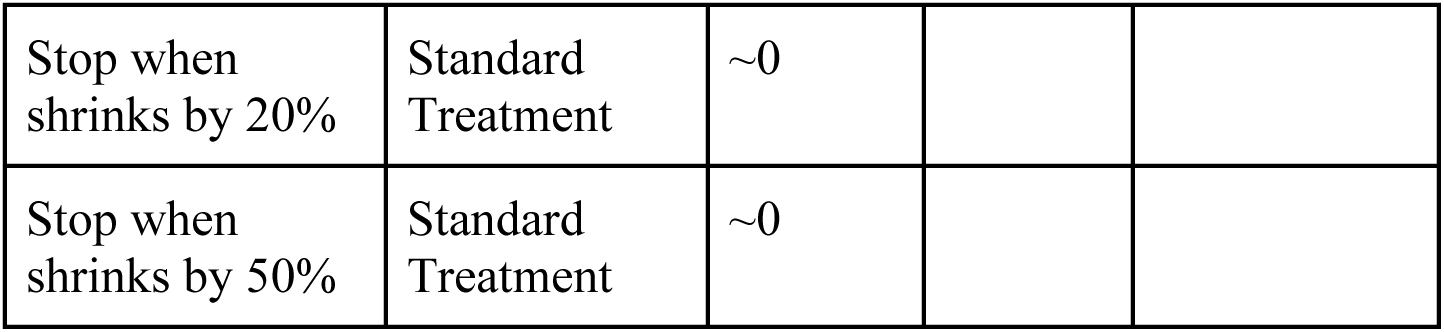
Effect of stopping treatment when tumor burden falls below a certain level for treatment using two cytotoxic or two cytostatic drugs.

**Table.8:**
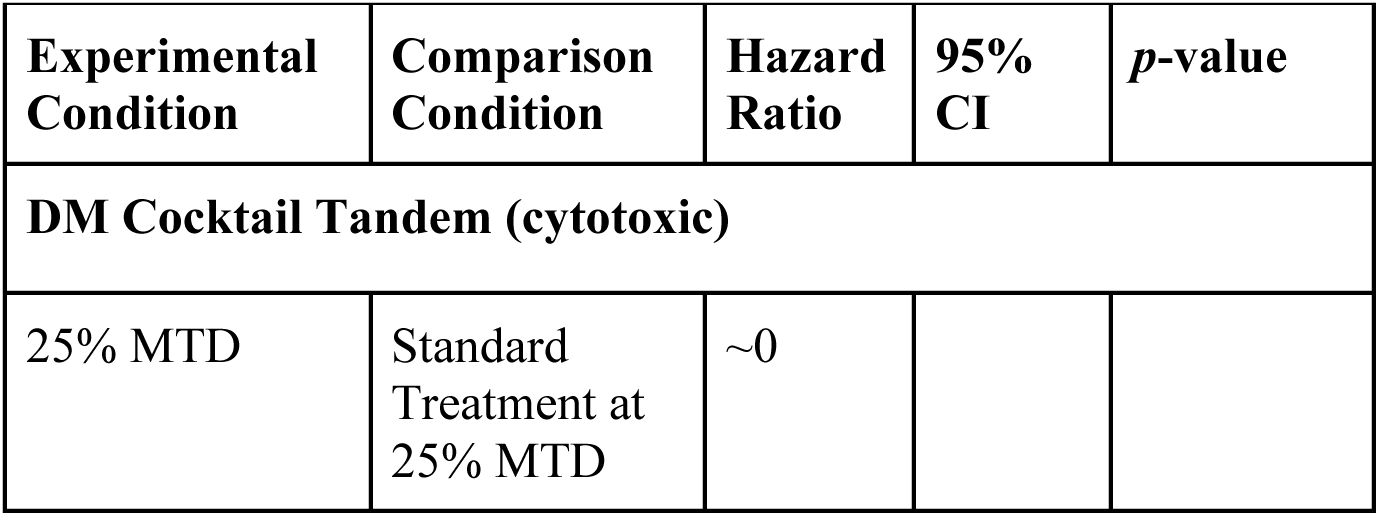
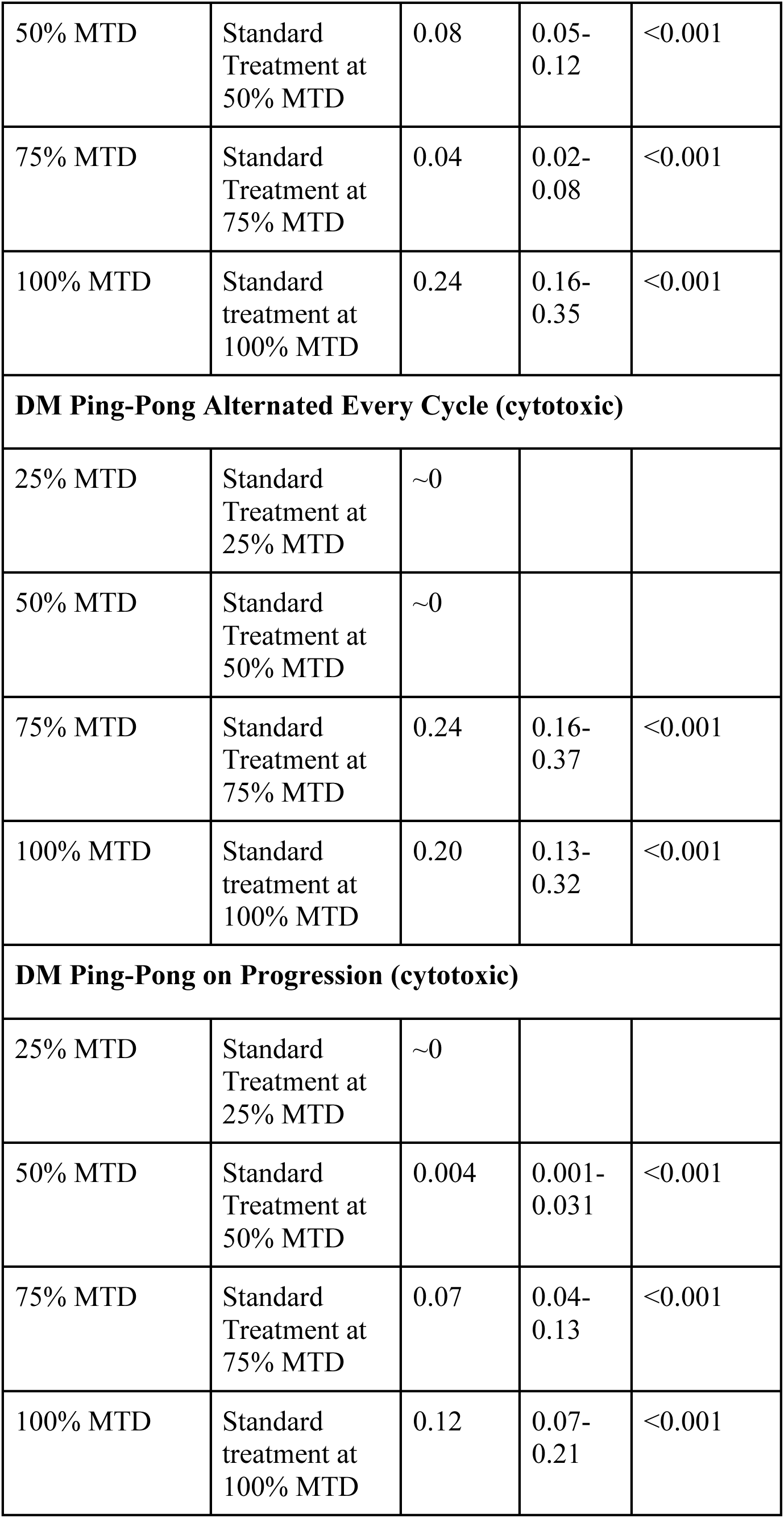
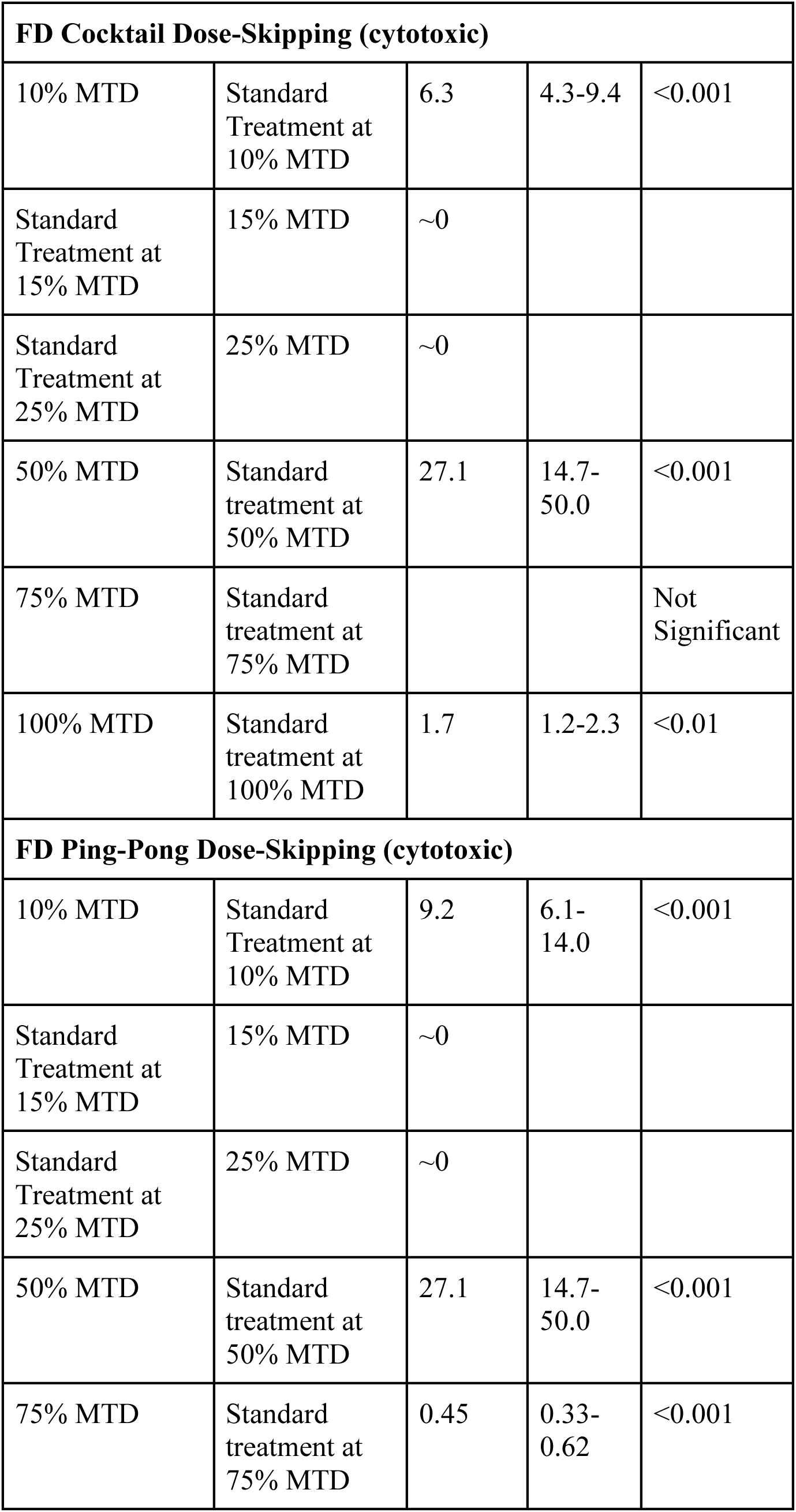
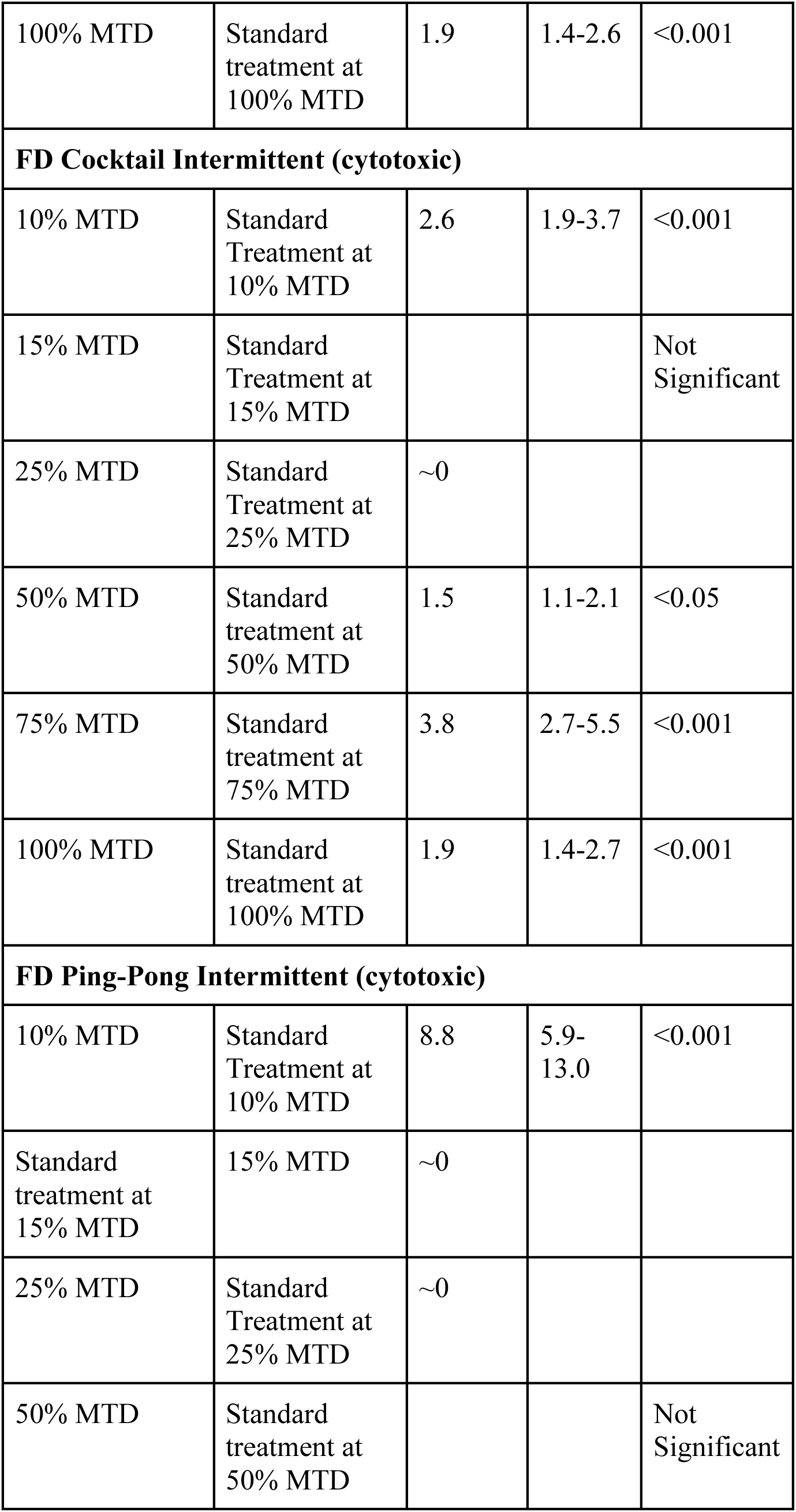
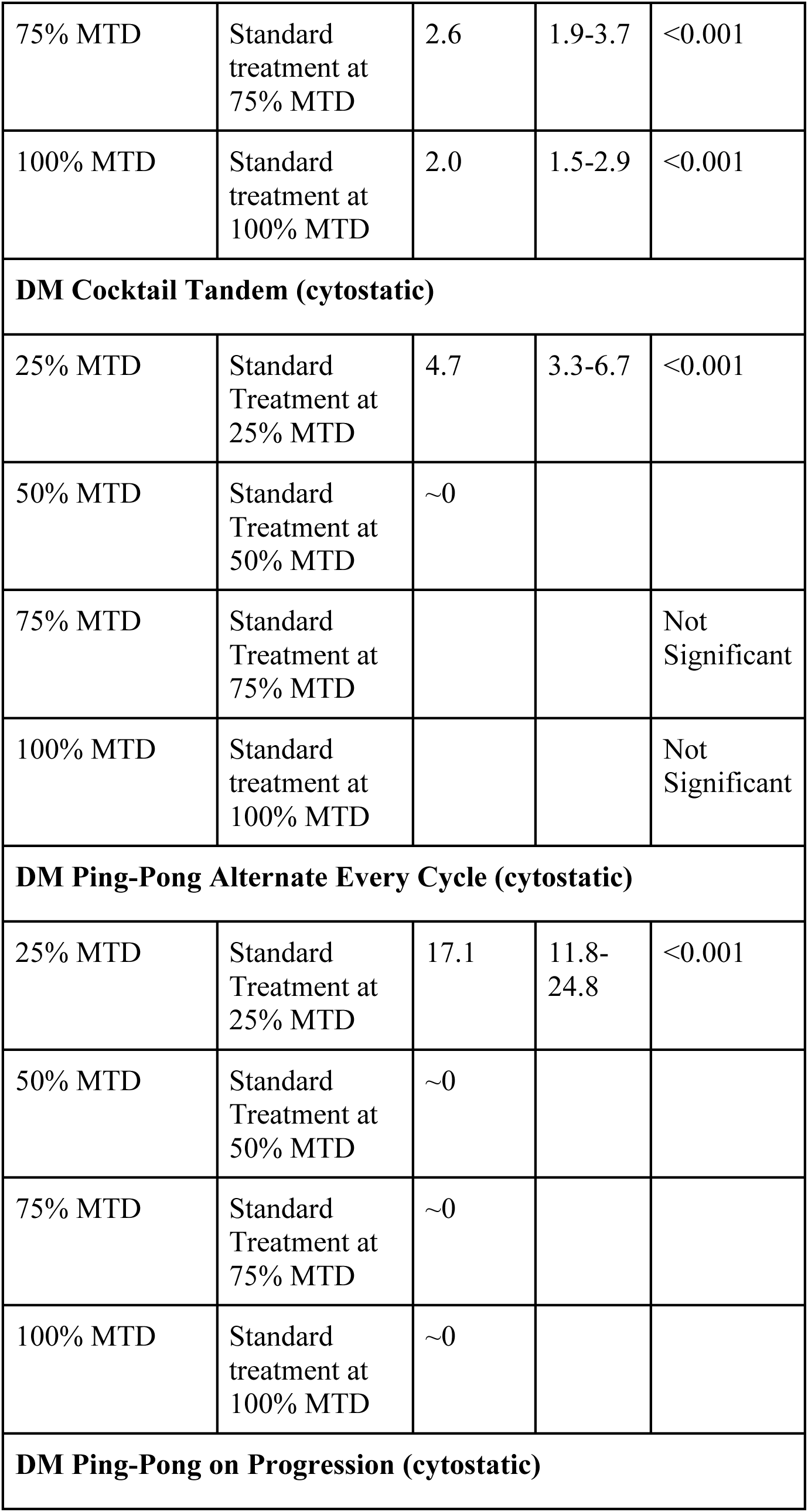
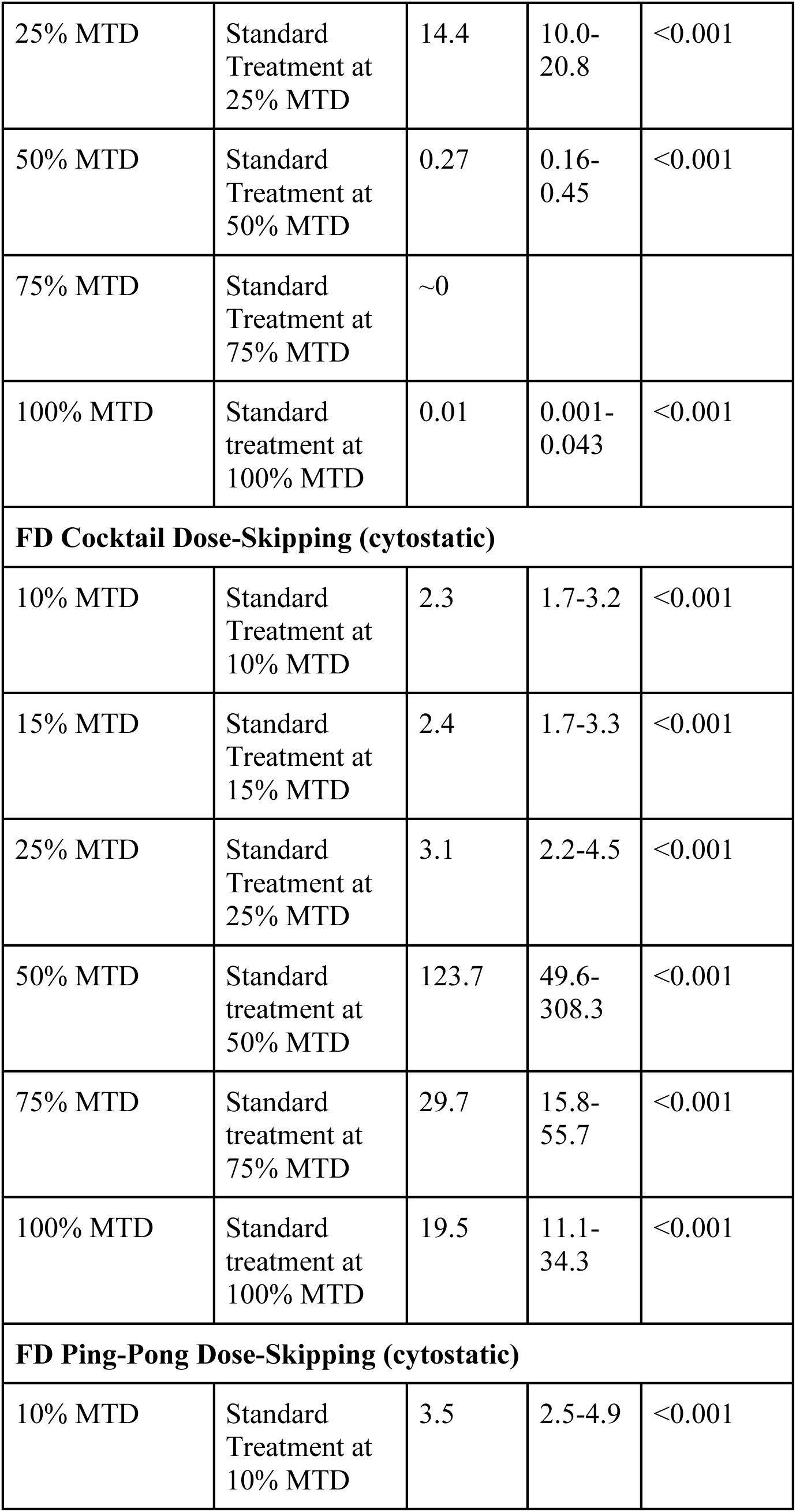
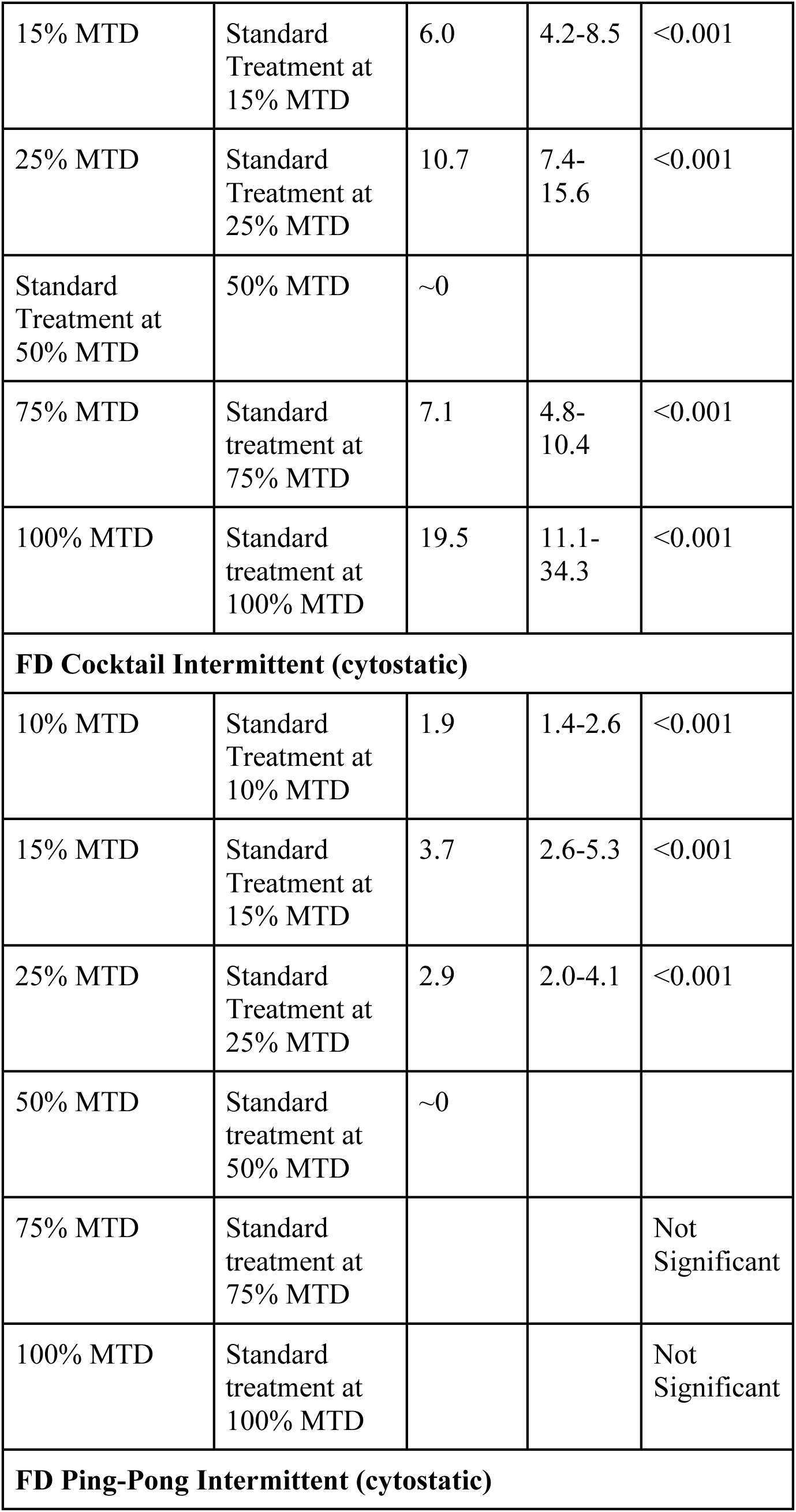
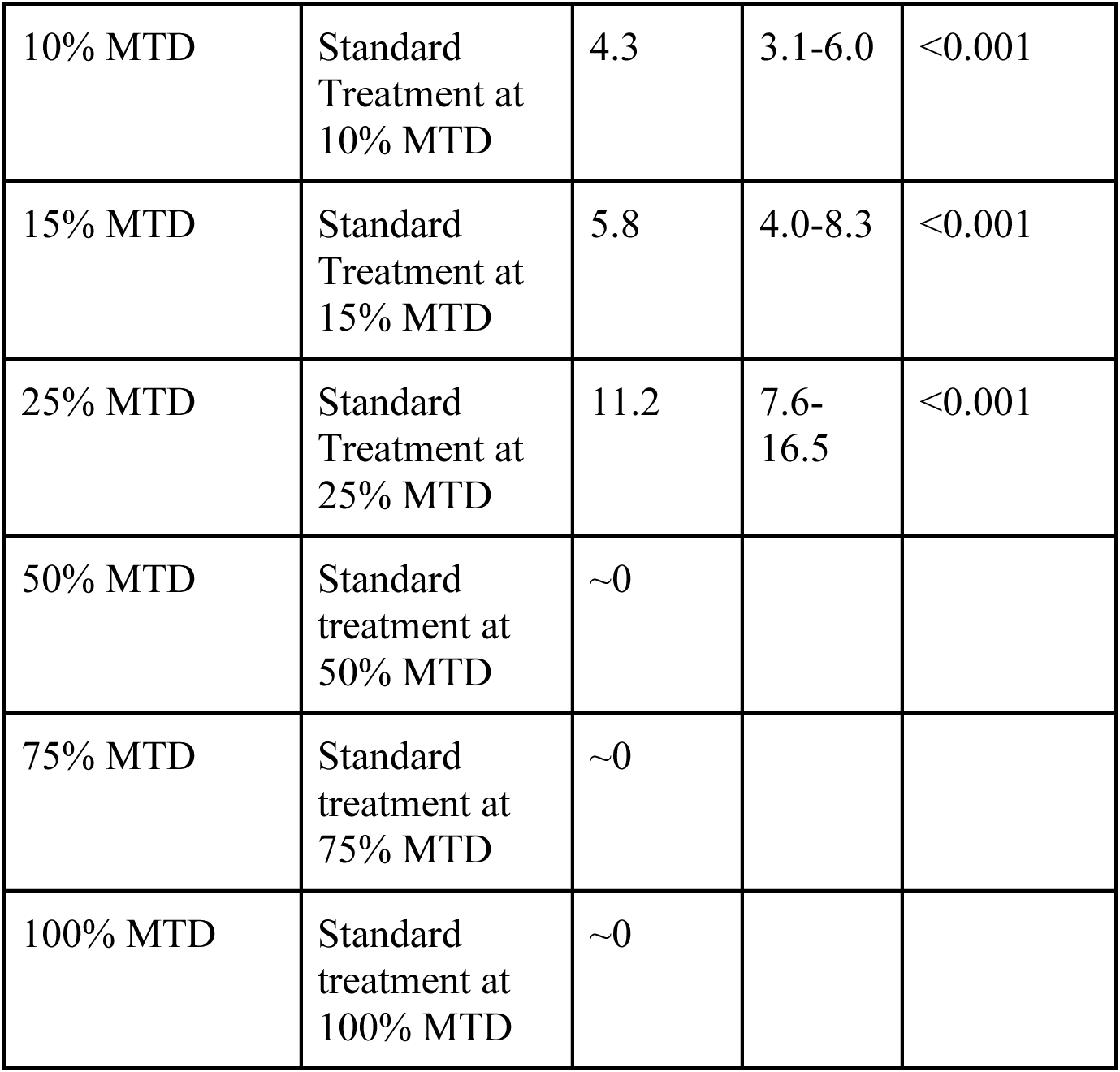
Effect of administering treatment at a range of different drug dosages for adaptive therapy using two cytotoxic or two cytostatic drugs.

| Experimental Condition | Comparison Condition | Hazard Ratio | 95% CI | p-value |
| --- | --- | --- | --- | --- |
| **DM Cocktail Tandem (cytotoxic)** |  |  |  |  |
| 25% MTD | Standard Treatment at 25% MTD | ~0 |  |  |
| 50% MTD | Standard Treatment at 50% MTD | 0.08 | 0.05-0.12 | <0.001 |
| 75% MTD | Standard Treatment at 75% MTD | 0.04 | 0.02-0.08 | <0.001 |
| 100% MTD | Standard treatment at 100% MTD | 0.24 | 0.16-0.35 | <0.001 |
| **DM Ping-Pong Alternated Every Cycle (cytotoxic)** |  |  |  |  |
| 25% MTD | Standard Treatment at 25% MTD | ~0 |  |  |
| 50% MTD | Standard Treatment at 50% MTD | ~0 |  |  |
| 75% MTD | Standard Treatment at 75% MTD | 0.24 | 0.16-0.37 | <0.001 |
| 100% MTD | Standard treatment at 100% MTD | 0.20 | 0.13-0.32 | <0.001 |
| **DM Ping-Pong on Progression (cytotoxic)** |  |  |  |  |
| 25% MTD | Standard Treatment at 25% MTD | ~0 |  |  |
| 50% MTD | Standard Treatment at 50% MTD | 0.004 | 0.001-0.031 | <0.001 |
| 75% MTD | Standard Treatment at 75% MTD | 0.07 | 0.04-0.13 | <0.001 |
| 100% MTD | Standard treatment at 100% MTD | 0.12 | 0.07-0.21 | <0.001 |

| **FD Cocktail Dose-Skipping (cytotoxic)** |  |  |  |  |
| --- | --- | --- | --- | --- |
| 10% MTD | Standard Treatment at 10% MTD | 6.3 | 4.3-9.4 | <0.001 |
| Standard Treatment at 15% MTD | 15% MTD | ~0 |  |  |
| Standard Treatment at 25% MTD | 25% MTD | ~0 |  |  |
| 50% MTD | Standard treatment at 50% MTD | 27.1 | 14.7-50.0 | <0.001 |
| 75% MTD | Standard treatment at 75% MTD |  |  | Not Significant |
| 100% MTD | Standard treatment at 100% MTD | 1.7 | 1.2-2.3 | <0.01 |
| **FD Ping-Pong Dose-Skipping (cytotoxic)** |  |  |  |  |
| 10% MTD | Standard Treatment at 10% MTD | 9.2 | 6.1-14.0 | <0.001 |
| Standard Treatment at 15% MTD | 15% MTD | ~0 |  |  |
| Standard Treatment at 25% MTD | 25% MTD | ~0 |  |  |
| 50% MTD | Standard treatment at 50% MTD | 27.1 | 14.7-50.0 | <0.001 |
| 75% MTD | Standard treatment at 75% MTD | 0.45 | 0.33-0.62 | <0.001 |
| 100% MTD | Standard treatment at 100% MTD | 1.9 | 1.4-2.6 | <0.001 |
| **FD Cocktail Intermittent (cytotoxic)** |  |  |  |  |
| 10% MTD | Standard Treatment at 10% MTD | 2.6 | 1.9-3.7 | <0.001 |
| 15% MTD | Standard Treatment at 15% MTD |  |  | Not Significant |
| 25% MTD | Standard Treatment at 25% MTD | ~0 |  |  |
| 50% MTD | Standard treatment at 50% MTD | 1.5 | 1.1-2.1 | <0.05 |
| 75% MTD | Standard treatment at 75% MTD | 3.8 | 2.7-5.5 | <0.001 |
| 100% MTD | Standard treatment at 100% MTD | 1.9 | 1.4-2.7 | <0.001 |
| **FD Ping-Pong Intermittent (cytotoxic)** |  |  |  |  |
| 10% MTD | Standard Treatment at 10% MTD | 8.8 | 5.9-13.0 | <0.001 |
| Standard treatment at 15% MTD | 15% MTD | ~0 |  |  |
| 25% MTD | Standard Treatment at 25% MTD | ~0 |  |  |
| 50% MTD | Standard treatment at 50% MTD |  |  | Not Significant |
| 75% MTD | Standard treatment at 75% MTD | 2.6 | 1.9-3.7 | <0.001 |
| 100% MTD | Standard treatment at 100% MTD | 2.0 | 1.5-2.9 | <0.001 |
| **DM Cocktail Tandem (cytostatic)** |  |  |  |  |
| 25% MTD | Standard Treatment at 25% MTD | 4.7 | 3.3-6.7 | <0.001 |
| 50% MTD | Standard Treatment at 50% MTD | ~0 |  |  |
| 75% MTD | Standard Treatment at 75% MTD |  |  | Not Significant |
| 100% MTD | Standard treatment at 100% MTD |  |  | Not Significant |
| **DM Ping-Pong Alternate Every Cycle (cytostatic)** |  |  |  |  |
| 25% MTD | Standard Treatment at 25% MTD | 17.1 | 11.8-24.8 | <0.001 |
| 50% MTD | Standard Treatment at 50% MTD | ~0 |  |  |
| 75% MTD | Standard Treatment at 75% MTD | ~0 |  |  |
| 100% MTD | Standard treatment at 100% MTD | ~0 |  |  |
| **DM Ping-Pong on Progression (cytostatic)** |  |  |  |  |
| 25% MTD | Standard Treatment at 25% MTD | 14.4 | 10.0-20.8 | <0.001 |
| 50% MTD | Standard Treatment at 50% MTD | 0.27 | 0.16-0.45 | <0.001 |
| 75% MTD | Standard Treatment at 75% MTD | ~0 |  |  |
| 100% MTD | Standard treatment at 100% MTD | 0.01 | 0.001-0.043 | <0.001 |
| **FD Cocktail Dose-Skipping (cytostatic)** |  |  |  |  |
| 10% MTD | Standard Treatment at 10% MTD | 2.3 | 1.7-3.2 | <0.001 |
| 15% MTD | Standard Treatment at 15% MTD | 2.4 | 1.7-3.3 | <0.001 |
| 25% MTD | Standard Treatment at 25% MTD | 3.1 | 2.2-4.5 | <0.001 |
| 50% MTD | Standard treatment at 50% MTD | 123.7 | 49.6-308.3 | <0.001 |
| 75% MTD | Standard treatment at 75% MTD | 29.7 | 15.8-55.7 | <0.001 |
| 100% MTD | Standard treatment at 100% MTD | 19.5 | 11.1-34.3 | <0.001 |
| **FD Ping-Pong Dose-Skipping (cytostatic)** |  |  |  |  |
| 10% MTD | Standard Treatment at 10% MTD | 3.5 | 2.5-4.9 | <0.001 |
| 15% MTD | Standard Treatment at 15% MTD | 6.0 | 4.2-8.5 | <0.001 |
| 25% MTD | Standard Treatment at 25% MTD | 10.7 | 7.4-15.6 | <0.001 |
| Standard Treatment at 50% MTD | 50% MTD | ~0 |  |  |
| 75% MTD | Standard treatment at 75% MTD | 7.1 | 4.8-10.4 | <0.001 |
| 100% MTD | Standard treatment at 100% MTD | 19.5 | 11.1-34.3 | <0.001 |
| **FD Cocktail Intermittent (cytostatic)** |  |  |  |  |
| 10% MTD | Standard Treatment at 10% MTD | 1.9 | 1.4-2.6 | <0.001 |
| 15% MTD | Standard Treatment at 15% MTD | 3.7 | 2.6-5.3 | <0.001 |
| 25% MTD | Standard Treatment at 25% MTD | 2.9 | 2.0-4.1 | <0.001 |
| 50% MTD | Standard treatment at 50% MTD | ~0 |  |  |
| 75% MTD | Standard treatment at 75% MTD |  |  | Not Significant |
| 100% MTD | Standard treatment at 100% MTD |  |  | Not Significant |
| **FD Ping-Pong Intermittent (cytostatic)** |  |  |  |  |
| 10% MTD | Standard Treatment at 10% MTD | 4.3 | 3.1-6.0 | <0.001 |
| 15% MTD | Standard Treatment at 15% MTD | 5.8 | 4.0-8.3 | <0.001 |
| 25% MTD | Standard Treatment at 25% MTD | 11.2 | 7.6-16.5 | <0.001 |
| 50% MTD | Standard treatment at 50% MTD | ~0 |  |  |
| 75% MTD | Standard treatment at 75% MTD | ~0 |  |  |
| 100% MTD | Standard treatment at 100% MTD | ~0 |  |  |

### Drug dosage level at which adaptive therapy is initiated and capped

Standard treatment for cytotoxic drugs begins to work at 25%of MTD as we lower the drug dosages, and works perfectly for 15% MTD, and at 10% begins to lose control of some of the tumors (Fig. 8G); for FD Cocktail Intermittent, 10% of MTD does not work but 15% and 25% work perfectly, and at 50% of MTD and above it does not work (Fig. 8G). Goldilocks effect in most of these cases is observed, where too much drug selects for resistance or too less drug is unable to control the tumor. At low extremes of the drug, the tumor grows out control, and at high extremes of the drug, the resistant clones are taking over the tumor. For cytostatic drugs, standard treatment at 25% or 50% of MTD works well (Fig. 8F), while standard treatment at 10%, 15%, 75%, and 100% performs poorly (Fig. 8F); for FD Cocktail Intermittent, treatment at 50% of MTD results in perfect survival, while survival outcome worsens with treatment at 10%, 15%, 25%, 75%, or 100% of MTD (Fig. 8H), exhibiting a Goldilocks effect for both standard treatment as well as adaptive therapy in many of these cases.

**Figure 8.**
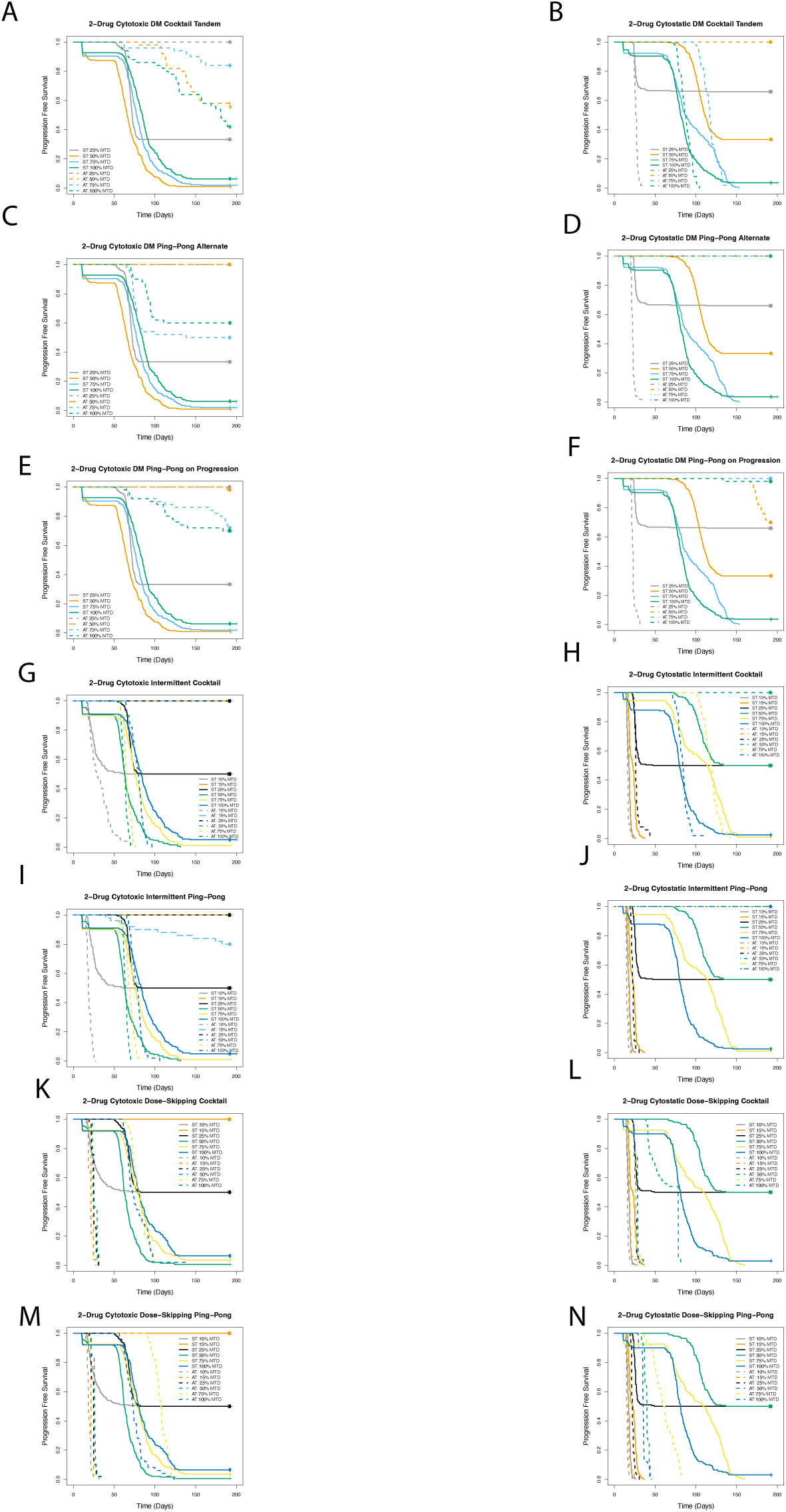
Effect of administering treatment at a range of different drug dosages for adaptive therapy using two cytotoxic or two cytostatic drugs. Survival outcome comparing treatment as per the dose modulation protocol relative to ST, starting and capping dosing at 25%, 50%, 75%, or 100% of MTD, for treatment using a single cytotoxic drug (Fig. 8A), or a single cytostatic drug (Fig. 8B). Survival outcome for treatment as per dose-skipping protocol administered at 35%, 50%, 75%, or 100% of MTD relative to standard treatment using either a single cytotoxic drug (Fig. 8C), or a single cytostatic drug (Fig. 8D). Survival outcome for treatment as per the intermittent protocol administered at 10%, 15%, 25%, 50%, 75%, or 100% of MTD relative to ST for treatment using a single cytotoxic (Fig. 8E), or a single cytostatic (Fig. 8F) drug.

### Median TTP versus Average Drug Dose

I plotted log10 median TTP versus average drug dose including all data points except for which a median TTP is not available (since less than 50% of the test subjects has progressed) and fitted the curve to a quadratic plot. For treatment using two cytotoxic drugs (Fig. 9A), median R-squared value is 0.03479 and adjusted R-squared is 0.0206. For treatment using two cytostatic drugs (Fig. 9B), median R-squared value is 0.6165 and adjusted R-squared value is 0.6105.

**Figure 9.**
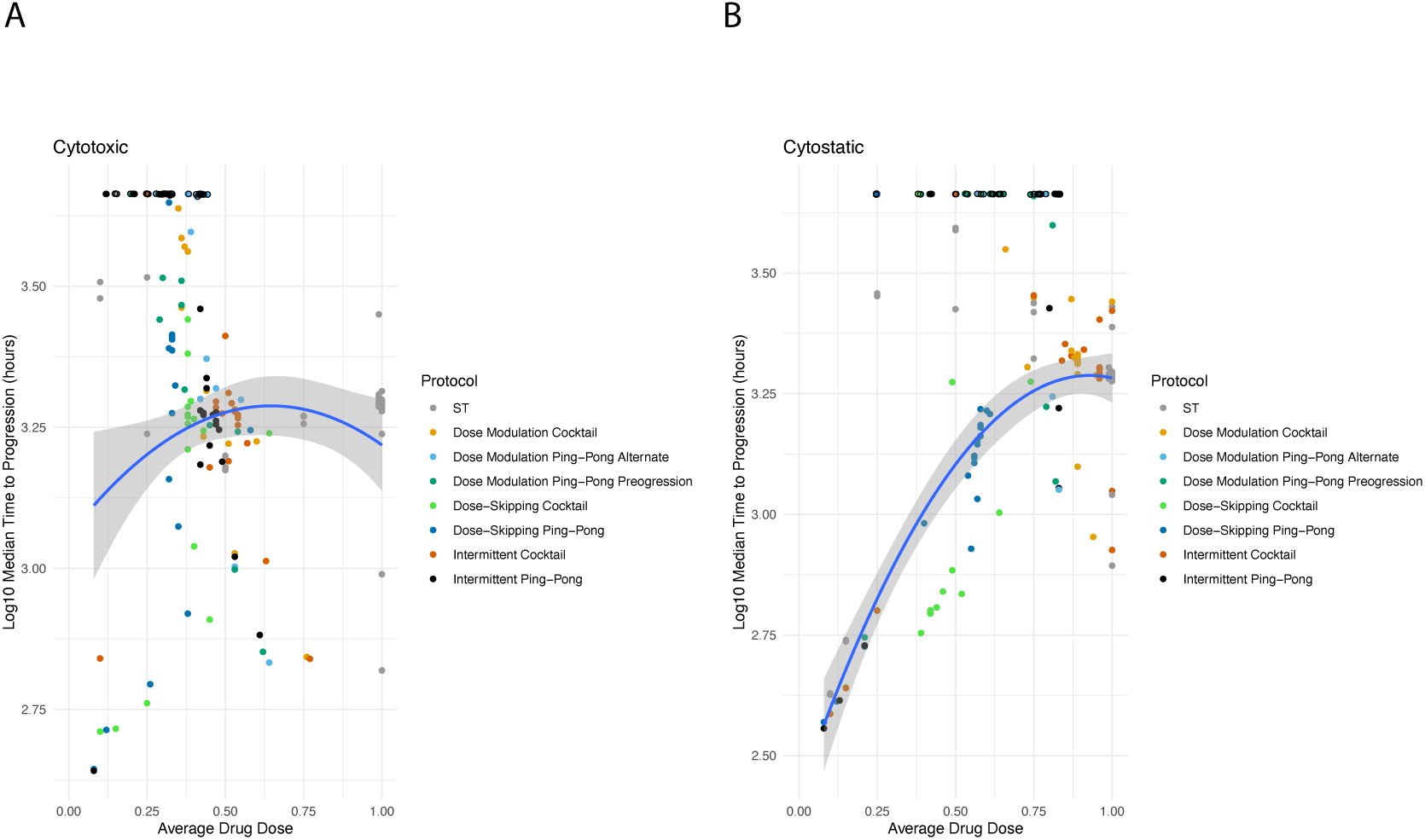
Summarizing the relationship between drug dose and time to progression for adaptive therapy using two cytotoxic or two cytostatic drugs. In each panel, the average amount of drug used per timestep between the start of therapy and the time of progression is plotted on the X-axis, and the median time to progression for that protocol under those parameter values are plotted on the Y-axis, for treatment using either a single cytotoxic drug (A) or a single cytostatic drug (B). The points are colored based on the specific protocol. Open circles indicate data points that are censored as less than 50% of test-subjects have progressed, and are not included in the calculation. A quadratic fit to the curve along with the confidence intervals have been indicated in the figure panels. Each point represents a specific Kaplan-Meier survival curve for a given set of parameter values.

## Discussion

Designing adaptive therapy protocols using two drugs is challenging as the number of parameters increase exponentially with each additional drug. Furthermore, we seek to investigate how the mechanism of action of the drugs, that is, whether or not the drugs are cytotoxic or cytostatic impacts survival outcome. Thus, we have investigated seven different treatment protocols, the dose modulation (DM) protocols: DM Cocktail Tandem, DM Ping-Pong Alternate Every Cycle, DM Ping-Pong on Progression; and the fixed-dose (FD) protocols: FD Cocktail Dose-Skipping, FD Ping-Pong Dose-Skipping, FD Cocktail Intermittent, FD Ping-Pong Intermittent; for treatment using either two cytotoxic or two cytostatic drugs.

We observed, for treatment using two cytotoxic drugs, all dose modulation protocols, namely, DM Cocktail Tandem, DM Ping-Pong Alternate Every Cycle, and DM Ping-Pong on Progression, as well the fixed-dose protocol FD Ping-Pong Dose-Skipping work well, increasing TTP relative to standard treatment. For treatment using two cytostatic drugs, the ping-pong protocols, namely, DM Ping-Pong Alternate Every Cycle, DM Ping-Pong on Progression, and FD Ping-Pong Intermittent work well, improving survival outcome relative to standard treatment.

Our results show that fitness cost, as manifested in increased doubling time for the drug-resistant cells in the absence of the drug, is required for adaptive therapy to work for both treatments using two cytotoxic or two cytostatic drugs. In general, a higher fitness cost leads to improved survival outcome, increasing TTP relative to lower fitness costs. However, a caveat with very high fitness cost is that the tumor burden is also increasing at a faster rate, which could translate to hitting the carrying capacity, and this effect might be especially pronounced under low amounts of the drugs. We observed FD Dose-Skipping treatment protocols working poorly under high fitness costs.

Cell replacement is an important parameter in our model as it is a parameter that governs whether or not a dividing cell would be allowed to replace a neighbor when there is no space available in the Moore neighborhood and, as such it is a measure of cell competition in the tumor. Our results indicate adaptive therapy works best at higher replacement rates and, in general, works best under conditions of 100% replacement. Furthermore, we observe this effect for both cytotoxic as well as cytostatic drugs. This suggests that adaptive therapy might work especially well when there is strong competition among the cell types constituting the tumor.

We observe, in general, adaptive therapy using two cytotoxic drugs works best under conditions of high turnover. However, in general, treatment using two cytostatic drugs seems to work best when the cell turnover is low.

We find, in general, adaptive therapy works best when drug doses are adjusted as soon as a change in tumor burden is detected, for treatment using either two cytotoxic or two cytostatic drugs. Furthermore, in general, the survival outcome worsens as Delta Tumor is progressively increased, with the best survival outcome with Delta Tumor=5% and the worst with Delta Tumor=40%.

We find, in general, for treatment using two cytotoxic or two cytostatic drugs, that it is best to pause treatment sooner than later when the tumor is shrinking. As such, for treatment as per the dose modulation protocol, triggering a treatment vacation when the tumor has shrunk by 20% works better than triggering a treatment vacation when the tumor has shrunk by 50%, which in turn works better than triggering a treatment vacation when the tumor has shrunk by 90%. We observe the same trend for treatment as per the intermittent protocol. As such, choosing to pause treatment when the tumor shrinks by 5% works better than choosing to pause treatment when the tumor shrinks by 10%, which is works better than choosing to pause treatment when the tumor shrinks by 20%, which work better than choosing to pause treatment when the tumor shrinks by 50%.

We observe, in general, for both treatment using either two cytotoxic or two cytostatic drugs, that too low dosage of a drug has poor survival outcome, and too high dosage of a drug also has poor survival outcome, with intermediate level of drug dosages having the best survival outcome. Thus, a Goldlilocks level of drug dosing works best for both treatments using either two cytotoxic or two cytostatic drugs. Consistent with this observation, fitting median TTP versus average drug dose to a quadratic results in significant fit.

There are several limitations to this work. We have not modeled tissue architecture or normal cells inside the tumors. Modeling blood vasculature and capillary architecture could affect treatment outcomes in important ways. In the future, it would be interesting to explore how these adaptive therapy protocols would perform in a 3-dimensional tumor model perfused with blood vessels and capillaries.

## Conclusions

While the dose modulation protocols (DM Cocktail Tandem, DM Ping-Pong Alternate Every Cycle, DM Ping-Pong on Progression), as well as the fixed-dose protocol (FD Ping-Pong Dose-Skipping) works well for treatment using two cytotoxic drugs, the ping-pong protocols except dose-skipping (DM Ping-Pong Alternate Every Cycle, DM Ping-Pong on Progression, and FD Ping-Pong Intermittent) works well for treatment using two cytostatic drugs, increasing TTP relative to standard treatment at maximum tolerated dose. In general, we observe adaptive therapy protocols work best under conditions of higher fitness cost, increased replacement rates, and higher turnover, with the exception of cytostatic drugs generally working better under conditions of low turnover. For the dose modulation protocols, as well as fixed-dose protocols, adaptive therapy works best when the doses are adjusted as soon as a change in tumor burden is detected. For the dose modulation protocols, in general, a relatively higher value for the delta dose parameter works best. We find that it is best to pause treatment sooner than later, when the tumor is shrinking. As such, triggering a treatment vacation or choosing to pause treatment when the tumor has shrunk by a relatively lower percentage relative to the baseline tumor burden at which therapy was initiated, works better than waiting for the tumor to shrink by a relatively higher percentage relative to that baseline. In general, for treatment using either two cytotoxic or two cytostatic drugs, an intermediate level of drug dosage works best, and worse survival outcome is observed with too little or too much drug, in accordance with the Goldilocks’ principle. Our results suggest adaptive therapy can be considerably improved by developing accurate and sensitive measurements of tumor burden, as well as by assaying for the level of cell turnover inside the tumor. Furthermore, our results lend support to the idea that it is indeed possible to transform cancer from an acute and lethal disease, which results in death, to a chronic disease, which does not lead to death.

